# Inference of coevolutionary dynamics and parameters from host and parasite polymorphism data of repeated experiments

**DOI:** 10.1101/625301

**Authors:** Hanna Märkle, Aurélien Tellier

## Abstract

There is a long-standing interest in understanding host-parasite coevolutionary dynamics and associated fitness effects. Increasing amounts of genomic data for both interacting species offer a promising source to identify candidate loci and to infer the main parameters of the past coevolutionary history. However, so far no method exists to do so. By coupling a gene-for-gene model with coalescent simulations, we first show that three types of biological cost, resistance, infectivity and infection, define the allele frequencies at the internal equilibrium point of the coevolution model, which in return determine the strength of the selective signatures signatures at host and parasite loci. We apply an Approximate Bayesian Computation (ABC) approach on simulated datasets to infer these costs by jointly integrating host and parasite polymorphism data at the coevolving loci. To control for the effect of genetic drift on coevolutionary dynamics, we assume that 10 or 30 repetitions are available from controlled experiments or several natural populations. We study two scenarios: 1) the cost of infection and population sizes (host and parasite) are unknown while costs of infectivity and resistance are known, and 2) all three costs are unknown while populations sizes are known. Using the ABC model choice procedure, we show that for both scenarios, we can distinguish with high accuracy pairs of loci from host and parasite under coevolution from neutrally evolving loci, though the statistical power decreases with higher cost of infection. The accuracy of parameter inference is also very high under both scenarios especially when using both host and parasite data because parasite polymorphism data do inform on host costs and vice-versa. As the false positive rate to detect genes under coevolution is small, we suggest to use our method to identify host and parasite candidate loci for further functional studies.

**Author summary:** It is of importance for agriculture and medicine to understand host-parasite antagonistic coevolutionary dynamics and the deleterious associated fitness effects, as well as to reveal the genes underpinning these interactions. The increasing amount of genomic data for hosts and parasites offer a promising source to identify such candidate loci, but also to use statistical inference methods to reconstruct the past coevolutionary history. In our study we attempt to draw inference of the past coevolutionary history at key host and parasites loci using sequence data from several individuals and across several experimental replicates. We demonstrate that using a Bayesian statistical method, it is possible to estimate the parameters driving the interaction of hosts and parasites at these loci for thousands of generations. The main parameter that can be estimated is the fitness loss by hosts upon infection. Our method and results can be applied to experimental coevolution data with sequences at the key candidate loci providing enough repetitions and large enough population sizes. As a proof of principle, our results open the door to reconstruct past coevolutionary dynamics using sequence data of interacting species.

## Introduction

Host-parasite coevolution is an ubiquitous process and has been demonstrated in terrestrial [1], limnological [2] and marine environments [3]. It describes the process of parasites and hosts exerting reciprocal selective pressures on one another. Therefore, coevolutionary dynamics are expected to substantially interact with and shape neutral nucleotide diversity linked to the coevolving sites. The latter can be single or multiple SNPs in coding or non-coding parts of genes [4,5], insertions/deletions [6] or distributed across a gene network [7]. Accordingly, the polymorphism patterns at the coevolving loci, referred to as the genetic signatures, are expected to be distinct from loci not involved into the coevolutionary interaction. Therefore, host and parasite genomic data are valuable source to identify loci under coevolution and to infer their past coevolutionary history.

On the one hand, signatures of positive selection which are characterized by lower genetic diversity compared to the genome-wide average and increased levels in linkage disequilibrium [8] are expected to arise under so called arms-race dynamics [9,10]. In arms race dynamics, frequencies of new beneficial alleles (such as new resistance or infectivity alleles) arising by *de novo* mutations increase towards fixation in both interacting partners. Accordingly, alleles are short lived and recurrently replaced and thus, allelic polymorphism is only transient [9,10]. On the other hand, signatures of balancing selection characterized by higher than average diversity [11] are expected to be the result of so called trench-warfare dynamics (also referred to as Red Queen dynamics) [6,9]. In this type of dynamics, several alleles are stably maintained over large time periods in both coevolving species. Hereby, allele frequencies either converge towards a stable equilbrium or they fluctuate persistently over time. Based on these classic expectations, genomic studies have unravelled positive and balancing selection signatures at various resistance genes [4–6,12–16] and effector genes [17,18]. An additional difficulty for coevolutionary analyses, is that there is a continuum between arms-race and trench-warfare dynamics and the dynamics are in fact strongly affected by the type and strength of various forms of selection (negative indirect frequency dependent selection, negative direct frequency dependent selection, overdominant selection) and their interplay with genetic drift [19,20] and mutation [21,22]. In other words, the expectations above are probably too simple to be accurately applicable as the effect of genetic drift and mutation affecting the outcome of coevolutionary dynamics is ignored [19–21].

Under negative frequency-dependent selection (nFDS) the fitness of a particular allele is either inversely proportional to its own frequency (direct, ndFDS) or allele frequencies in the interacting partner (indirect, niFDS) [23,24]. Overdominant selection or some form of ndFDS are a necessary but not always sufficient condition for trench-warfare dynamics to take place in single locus host-parasite coevolutionary interactions [21,24]. But even with some form of ndFDS acting, arms-race dynamics can take place if either the strength of ndFDS compared to niFDS is weak or genetic drift is causing random loss of alleles.

The exact nature of the dynamics, such as the equilibrium frequencies of alleles and the period and amplitude of coevolutionary cycles, is further affected by the way host and parasite genotypes interact at the molecular level and the fitness costs associated with the coevolutionary interaction. The interaction at the molecular level is captured by the infection matrix which stores the specificity and level of infection in all possible pairwise interactions between host and parasite genotypes [25]. One well studied type of interaction is the gene-for-gene (GFG) interaction which presents one endpoint of a continuum of infection matrices [26,27]. GFG-interactions are characterized by one universally infective parasite genotype and one universally susceptible host type and for example have been found in the Flax-*Melampsora lini* system [28].

A fitness cost which has been shown to crucially affect the coevolutionary dynamics is the loss in host fitness due to infection [19,24]. In addition, costs of resistance such as reduced competitive ability or fertility in absence of the parasite [29–31] and costs of infectivity such as reduced spore production of infective pathogens [32] can further alter the dynamics. These costs also determine the equilibrium frequencies of the coevolutionary system [33,34] at which one or several alleles are maintained or around which allele frequencies cycle. An important result from previous theoretical investigations [33,34] is that the equilibrium frequencies in the parasite population depend on the parasite fitness costs (cost of infectivity) and vice versa (cost of resistance and cost of infection).

Given this continuum of coevolutionary dynamics, it is necessary to gain a deeper and refined understanding on how the interaction between allele frequency dynamics at the coevolving loci, genetic drift and mutation shapes the resulting genetic signatures at the coevolutionary loci and linked neutral sites. This is an important step for the development and application of methods designed to draw inference on the coevolutionary history. A previous study has investigated this link for two distinct coevolutionary models [19]. Focusing on a small set of summary statistics, the signatures at the coevolving loci cannot be necessarily distinguished from neutrality when considering host or parasite data in isolation. Moreover, the strength of coevolutionary signatures depends on the host and parasite population sizes and varies with changing costs of infection, resistance and infectivity [19].

The first aim of the present paper is to extend this approach [19] by including additional summary statistics in order to get a more refined understanding of the resulting genetic signatures. Based on this extended set of summary statistics, our major aim is to jointly infer several of the above mentioned parameters as a proof-of-principle by using an Approximate Bayesian Computation approach [35–37]. We base our inference on average summary statistics from *r* = 10 and *r* = 30 repeatedly simulated coevolutionary histories. We test this approach on two different scenarios. In scenario 1, we aim to infer simultaneously the cost of infection (*s*), the host population size (*N*_*H*_) and the parasite population size (*N*_*P*_) assuming that we know the true cost of resistance (*c*_*H*_) and the true cost of infectivity (*c*_*P*_). This scenario mimics systems where experimental measures of the costs of resistance or infectivity have been performed [32,38] and thus, these parameters can be assumed as known. In scenario 2, our goal is to infer simultaneously the cost of infection (*s*), the cost of infectivity (*c*_*P*_) and the cost of resistance (*c*_*H*_) that the true host (*N*_*H*_) and parasite population sizes (*N*_*P*_) are known. Scenario 2 is motivated by the assumption that an independent estimate of the effective population size can be obtained by using full-genome data of loci unlinked to the coevolutionary locus. For each scenario we perform the ABC model choice to distinguish coevolutionary from neutral loci and subsequently infer the model parameters.

## Results

### Link between coevolutionary dynamics and sequence data

Previous work has dealt with understanding the coevolutionary dynamics under the chosen coevolution model [24] and the resulting genetic signatures [19]. We provide a short summary of these results here to help the reader to gain an intuition regarding the ABC results. A classic coevolutionary cycle in this Gene-For-Gene model consists of four phases (see S1 Fig, [24,53]):

1. The frequency of *RES* hosts increases when *INF*-parasites are in low frequency.
2. As a response to the increasing frequency of *RES*-host the frequency of *INF*-parasites increases very quickly and the parasite population reaches almost fixation for the *INF*-allele.
3. This results in a decrease of the frequency of *RES*-hosts due to the cost of resistance.
4. Once *RES*-hosts are in low frequency the frequency of *ninf*-parasites increases due to the cost of infectivity.

Depending on the combination of cost of infection (*s*), cost of resistance (*c*_*H*_) and infectivity (*c*_*P*_) the model either exhibits trench-warfare dynamics or arms-race dynamics. Trench-warfare dynamics mainly take place for small to intermediate costs of infection. the dynamics switch to arms-race for high costs of infection (S1 Fig, S2 Fig), irrespective of *c*_*H*_ and *c*_*P*_ When arms-race dynamics take place the parasite population always exhibits fixation of the *INF*-allele. The speed of the subsequent fixation of the *res*-allele in the host depends on the cost of resistance (*c*_*H*_) and is faster for higher costs of resistance (*c*_*H*_).

The internal equilibrium frequencies under trench-warfare dynamics are affected as follows. The frequency of *RES*-hosts mainly increases with increasing cost of infectivity (*c*_*P*_) (S2 Fig a+b vs. S2 Fig c+d), increases very slightly with increasing cost of infection (*s*) and remains almost unaffected by changing costs of resistance (*c*_*H*_) (S2 Fig a+c vs. S2 Fig b+d). The opposite is true for the parasite. Here, the equilibrium frequency of the infective (*INF*)-parasite rises mainly with increasing cost of infection (*s*) (S2 Fig). Higher costs of resistance (*c*_*H*_) decrease the equilibrium frequency of *INF*-parasites (S2 Fig a+c vs. S2 Fig b+d) for a given value of *s*. In contrast to the host, the equilibrium frequencies in the parasite are almost unaffected by changes in the cost of infectivity (*c*_*P*_).

The changes in equilibrium frequencies with changing cost of infection (*s*), cost of resistance (*c*_*H*_) and changing cost of infectivity (*c*_*P*_) are reflected by the resulting genetic signatures at the coevolving loci (S9 Fig). We summarize the genetic signatures of coevolution chiefly as the behaviour of Tajima’s D under selective sweeps and balancing selection (S9 Fig,S3 Fig) is well known. Generally, the strongest signatures of balancing selection, indicated by high Tajima’s D values, can be observed when the equilibrium frequencies of *INF*-parasites or *RES*-hosts are close to 0.5 (see S2 Fig, S9 Fig). The strength of the signatures declines the further the equilibrium frequencies move away from 0.5.

The genetic signature in the parasite changes strongly with changing cost of infection (*s*), irrespectively of *c*_*H*_ and *c*_*P*_ (S9 Fig). Further, the resulting genetic signatures in the parasites for a given cost of infection *s* are distinguishable for different costs of resistance but not for different costs of infectivity. The genetic signature in the host is mainly indicative about the cost of infectivity (*c*_*P*_), a cost which is affecting the parasite fitness, whereas the signature in the parasite is mainly informative about the costs of resistance (*c*_*H*_) and infection (*s*), parameters with a direct fitness effect on the host (S9 Fig). The qualitative changes of the genetic signatures for changing costs of infection remain similar even when population sizes differ in both interacting partners (S3 Fig). However, their strength is affected by the population sizes.

### Inference of coevolutionary dynamics from polymorphism data

#### Scenario 1

##### Model choice

Under scenario 1 and *r* = 30, the model choice procedure is suited to distinguish a coevolutionary model in which the cost of infection (*s*), host population size (*N*_*H*_) and parasite population size (*N*_*P*_) are unknown from a neutral model where the host and parasite population size are unknown. The cross-validation reveals (values for *r* = 10 in brackets) that 482 (441) out of 500 coevolution simulations are correctly identified, while 18 (59) are misclassified as neutrally evolving pairs of loci, yielding a FNR of 3.6% (respectively 11.8%). In addition, 498 (495) neutral simulated pairs of loci are correctly identified as evolving neutrally, yielding a FPR of 0.4% (respectively 1% for *r* = 10) (see S4 Fig, S5 Fig). When analysing results for the PODs the accuracy of model choice is very high for low costs of infection but becomes worst when *s* > 0.6 (Fig 1, S6 Fig). It is apparent from Fig 2 that for Tajima’s D and PMD all PODs with intermediate to high *s* are in the cloud of neutral simulations (see S7 Fig for *r* = 10). For high values of *s*, dynamics are indeed switching to arms-race generating fast recurrent selective sweeps. Hence, the values of these statistics become similar to neutral expectations under small host and parasite population sizes.

**Fig 1.**
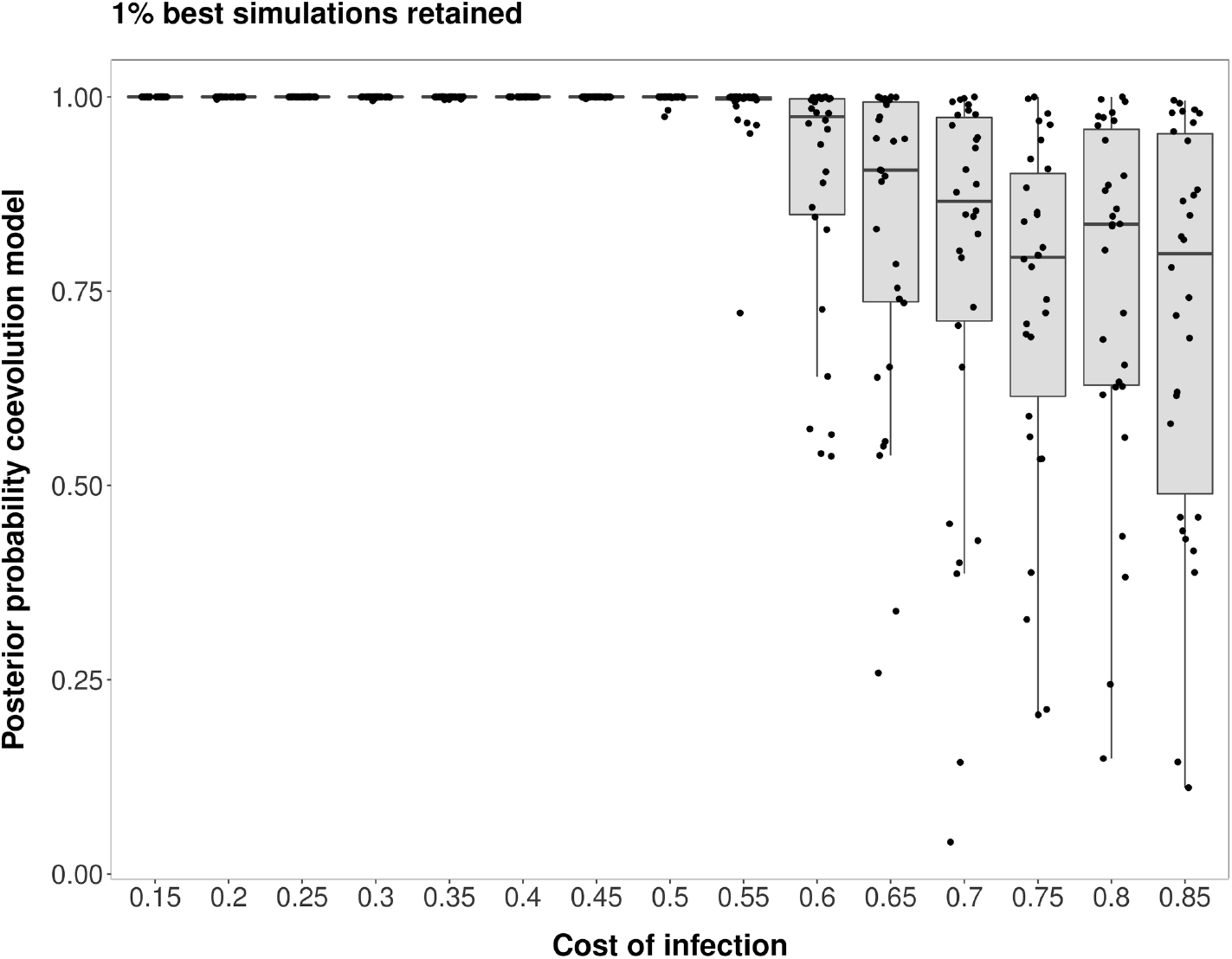
Posterior probability in support of the coevolution model (against a neutral model) for scenario 1. Results shown for 30 repetitions and 30 PODs per value of the cost of infection (*s*). Results for single PODs are shown as dots. Model choice distinguishing a coevolution model with unknown costs of infection (*s*), host population size (*N*_*H*_) and parasite population size (*N*_*P*_) from a neutral model with unknown host and parasite population sizes. Note that for these points we added some jitter to the x-values in order to increase the readability of the plots.

**Fig 2.**
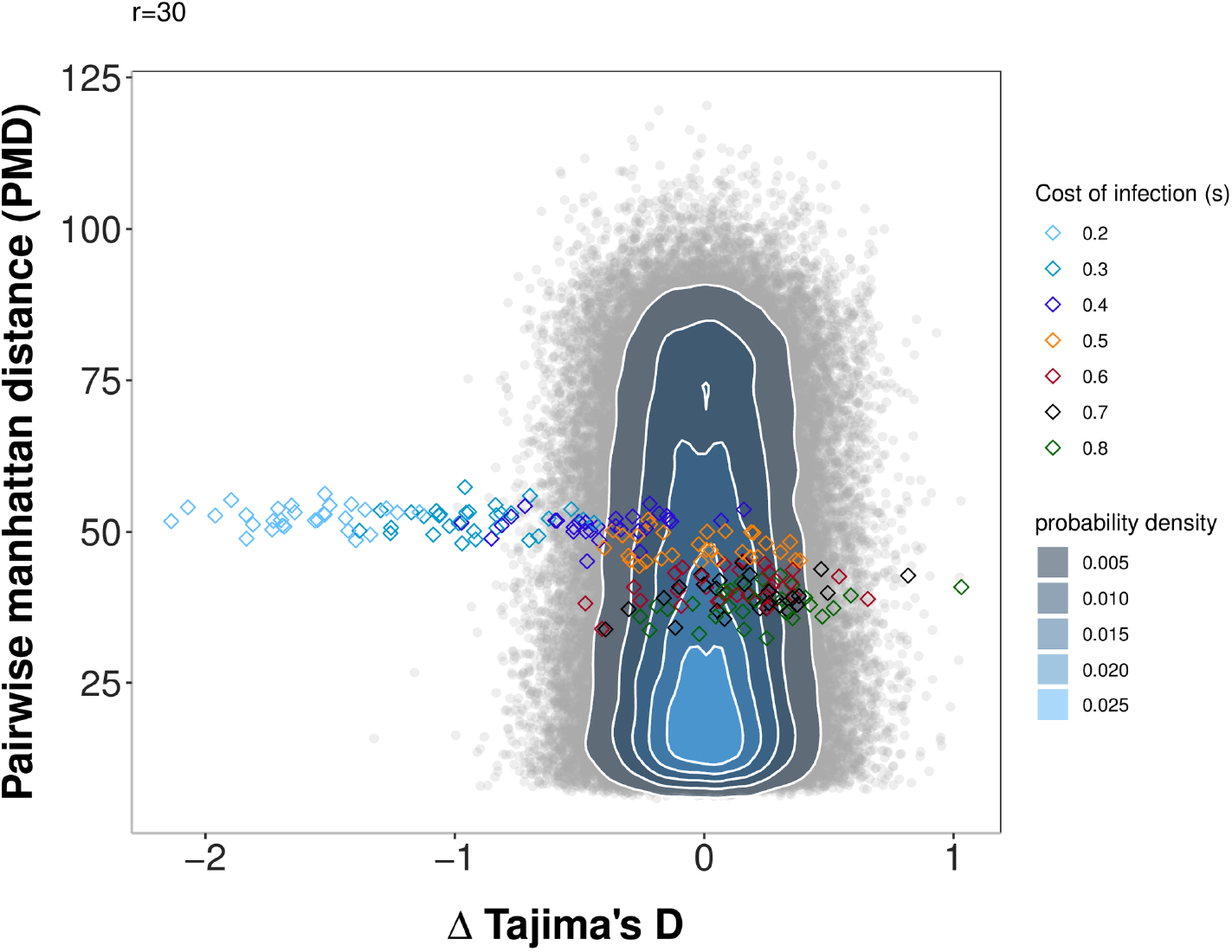
Pairwise Manhattan distance and Δ Tajima’s D (host-parasite) for the PODs under scenario 1 compared to simulations under a neutral model. Pairwise Manhattan distance (x-axis) and the difference between Tajima’s D of the host and of the parasite (y-axis) for the PODs used for inference in Scenario 1 and the 100,000 neutral simulations run for this scenario. Under the neutral model, host and parasite population sizes vary. Simulations under the neutral model are shown as grey open circles, and a bivariate normal kernel estimation has been applied to obtain a probability density of the summary statistic combinations. The PODs for scenario 1 are shown as diamonds and are coloured coded based on the true cost of infection (*s*).

##### Parameter estimation

Our results indicate that it is possible to jointly infer the cost of infection (*s*), the host population size (*N*_*H*_) and the parasite population size (*N*_*P*_) using polymorphism data from the host and parasite (Fig 3, S8 Fig). Generally, the accuracy of inference mainly depends on 1) the true value of the cost of infection and the 2) the type of polymorphism data being used (host and parasite together, only host or only parasite).

**Fig 3.**
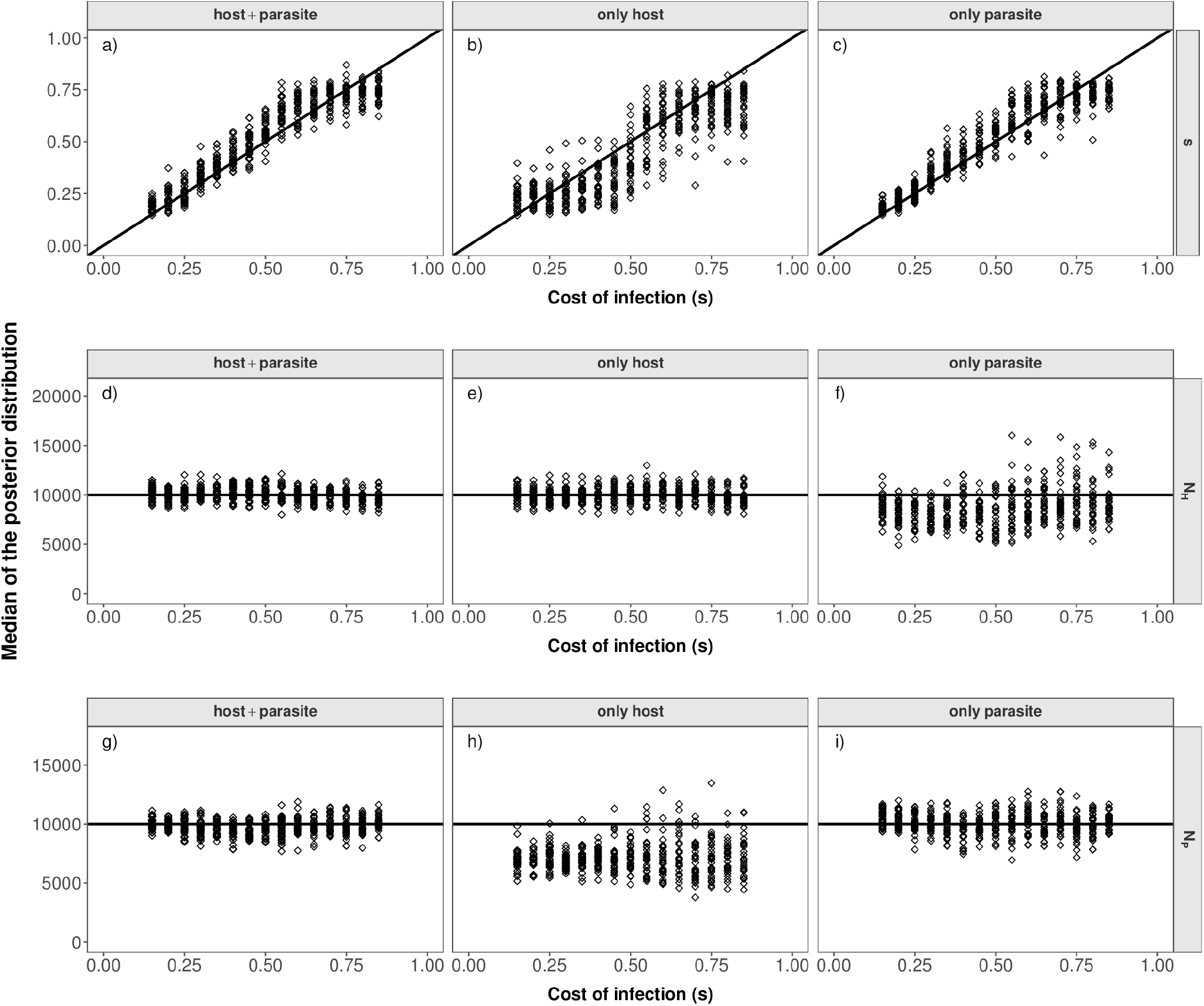
Parameter estimations under scenario 1. Median of the posterior distribution (y-axis) for the cost of infection *s* (top, a-c), host population size (*N*_*H*_) (middle, d-f) and parasite population size (*N*_*P*_) (bottom, g-i) when inference is based on host and parasite summary statistics (left), only host summary statistics (middle) or only parasite summary statistics (right). The median of the posterior distribution (after post-rejection adjustment) is plotted for each POD. The true cost of infection for each POD is shown on the x-axis.

Inferences of the cost of infection and of the population sizes are the most accurate if both host and parasite polymorphism data are used (Fig 3, S8 Fig). Using only parasite polymorphism data is also quite accurate for inferring small to intermediate values of the cost of infection (*s <* 0.6) (Fig 3 c+f) where trench-warfare dynamics take place. In contrast, using only host polymorphism data shows markedly less accuracy in the same parameter range (Fig 3 b+e). Overall the inference results become less accurate when decreasing the number of repetitions to *r* = 10 (S8 Fig).

#### Scenario 2

##### Model choice

Under scenario 2 and *r* = 30, model choice is suited to discriminate between coevolution and neutral evolution. Out of the 500 coevolution validation simulations 470 (417 for *r* = 10) are correctly classified as coevolving pairs of loci whereas 30 (83) are classified as neutrally evolving pairs, yielding a FNR of 6% (respectively 16.6% for *r* = 10). In addition, 489 (495 for *r* = 10) neutral simulated pairs of loci are correctly identified as evolving neutrally, yielding a FPR of 2.2% (respectively 1% for *r* = 10) (see S10 Fig, S11 Fig). When analysing results for the PODs the accuracy of model choice is very high under higher cost of infectivity (*c*_*P*_ = 0.3). For a lower value of *c*_*P*_ = 0.1, the model choice becomes less accurate when *s* increases, especially when *c*_*H*_ = 0.1 (Fig 4). As for scenario 1, it is also apparent from Fig 5 that some PODs are found within the cloud of neutral simulations (see S13 Fig for *r* = 10). For high values of *s*, dynamics are indeed switching to arms-race generating fast recurrent selective sweeps. Note however, the interesting case of *c*_*H*_ = *c*_*P*_ = 0.1 which displays the worst accuracy for high values of *s*. This is explained by very fast recurrent selective sweeps along with very fast coevolutionary cycles due to the combination of high cost of resistance and low cost of infectivity. Note that such fast cycles affect more strongly other statistics (in particular the nucleotide diversity) than the three we present in Figure 5 (Tajima’s D host and parasite and PMD), thus highlighting the need to include a larger number of summary statistics in the ABC procedure.

**Fig 4.**
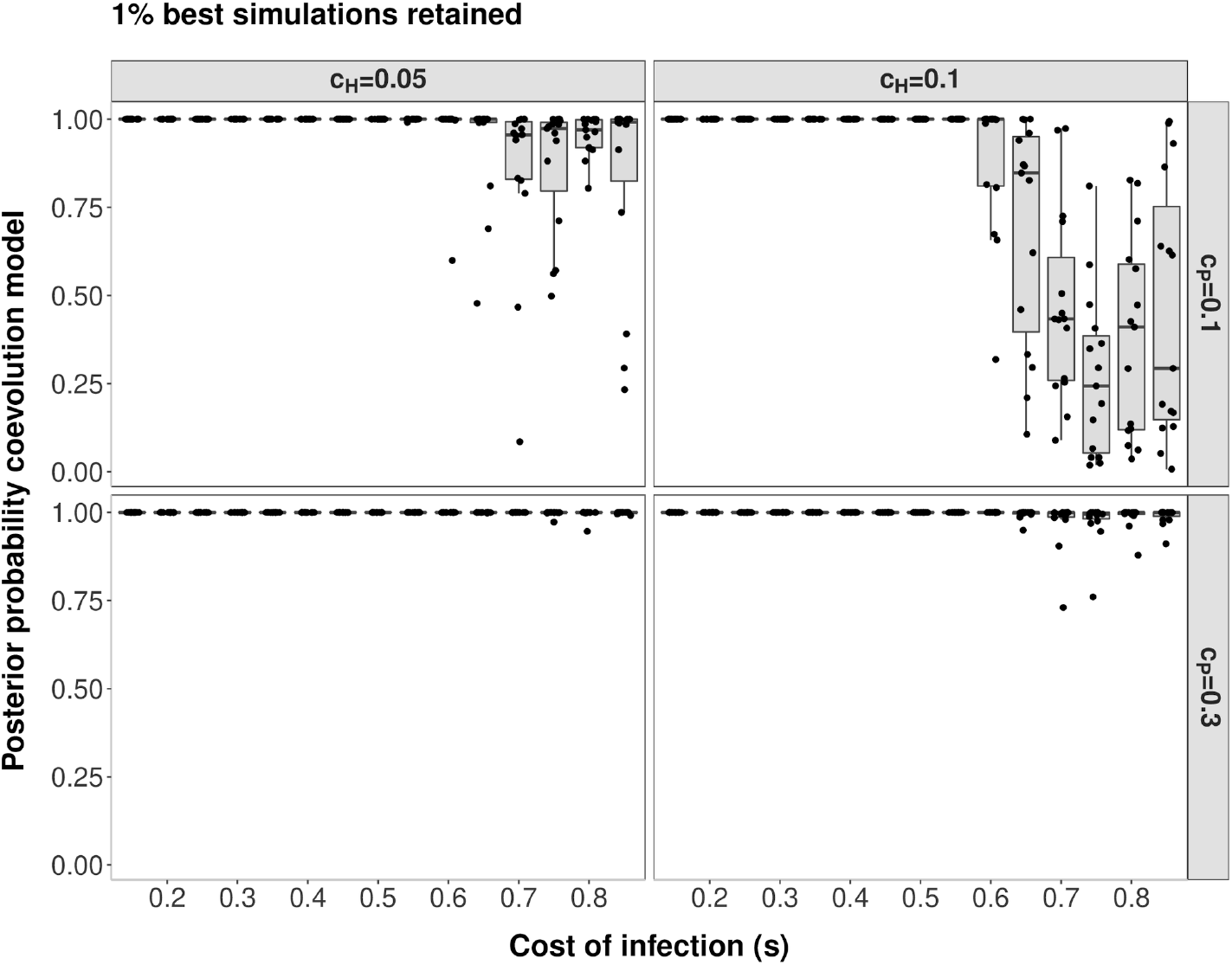
Posterior probability in support of the coevolution model (against a neutral model) for scenario 2. Results are shown for *r* = 30 and 15 PODs per boxplot. The posterior density in support of the coevolution model (y-axis) is shown for PODs with varying cost of infection (*s*). The different panels reflect the combination of *c*_*H*_ and *c*_*P*_ for the respective PODs (left: *c*_*H*_ = 0.05, right: *c*_*H*_ = 0.1, top: *c_P_* = 0.1, bottom: *c*_*P*_ = 0.3). Model choice has been run to distinguish a coevolution model with unknown costs of infection (*s*), cost of resistance (*c*_*H*_) and cost of infectivity (*c*_*P*_) from a neutral model with constant host and parasite population size (*N*_*H*_ = *N*_*P*_ = 10,000). Results for single PODs are shown as dots and jitter added to the x-values to increase the readability.

**Fig 5.**
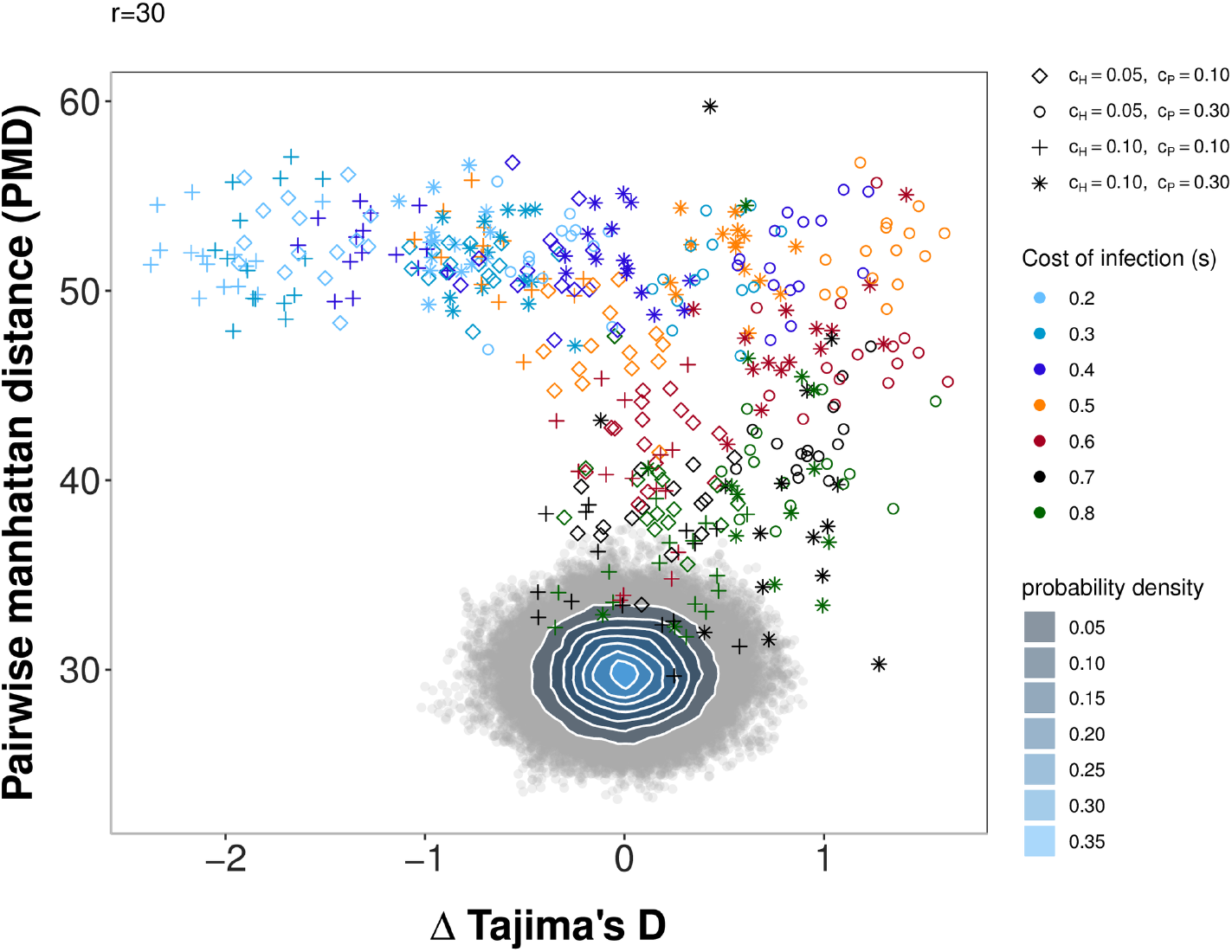
Pairwise Manhattan distance and Δ Tajima’s D (host-parasite) for the PODs under scenario 2 compared to simulations under a neutral model. Pairwise Manhattan distance (x-axis) and the difference between Tajima’s D of the host and of the parasite (y-axis) for the PODs used for inference in Scenario 2 and 100,000 neutral simulations. Simulations under the neutral model are shown as grey open circles. A bivariate normal kernel estimation has been applied to obtain a probability density of the different summary statistic combinations. The PODs for scenario 2 are shown in color. Colors reflect the true cost of infection (*s*) for a particular POD (see legend) and shapes indicate the combination of *c*_*H*_ and *c*_*P*_ (diamonds: *c*_*H*_ = 0.05, *c*_*P*_ = 0.1; circles: *c*_*H*_ = 0.05, *c*_*P*_ = 0.3; crosses: *c*_*H*_ = 0.01, *c*_*P*_ = 0.1; stars: *c*_*H*_ = 0.1, *c*_*P*_ = 0.3) for the respective POD.

##### Parameter estimation

As for scenario 1, the accuracy of inference for scenario 2 is best if data from both the host and the parasite are available (6). However, inference of the cost of infection *s* is less accurate compared to scenario 1. The most accurate inference results are obtained for intermediate costs of infection. This is due to the fact that signatures in the host and the parasite are differentially affected by the various costs (S9 Fig).

Inference of the cost of resistance (*c*_*H*_) works reasonably well if polymorphism data only from the parasite are available. However, this comes at the cost of less accurate inference of the cost of infection (*s*) as both parameters are affecting the equilibrium frequency in the parasite (S2 Fig, Fig 6). This effect is especially pronounced when the cost of infection (*s*) is low and only the information from the parasite polymorphism data are available (see Fig 6 c+f). The inference of the cost of infectivity (*c*_*P*_) is reasonably accurate if polymorphism data only from the host are available (Fig 6 h+q). This is due to the fact that the cost of infectivity (*c*_*P*_) mainly affects the equilibrium frequency in the host but not in the parasite (S2 Fig). Therefore, inference of this parameter does not work if only parasite polymorphism data are available (Fig 6 i+r).

**Fig 6.**
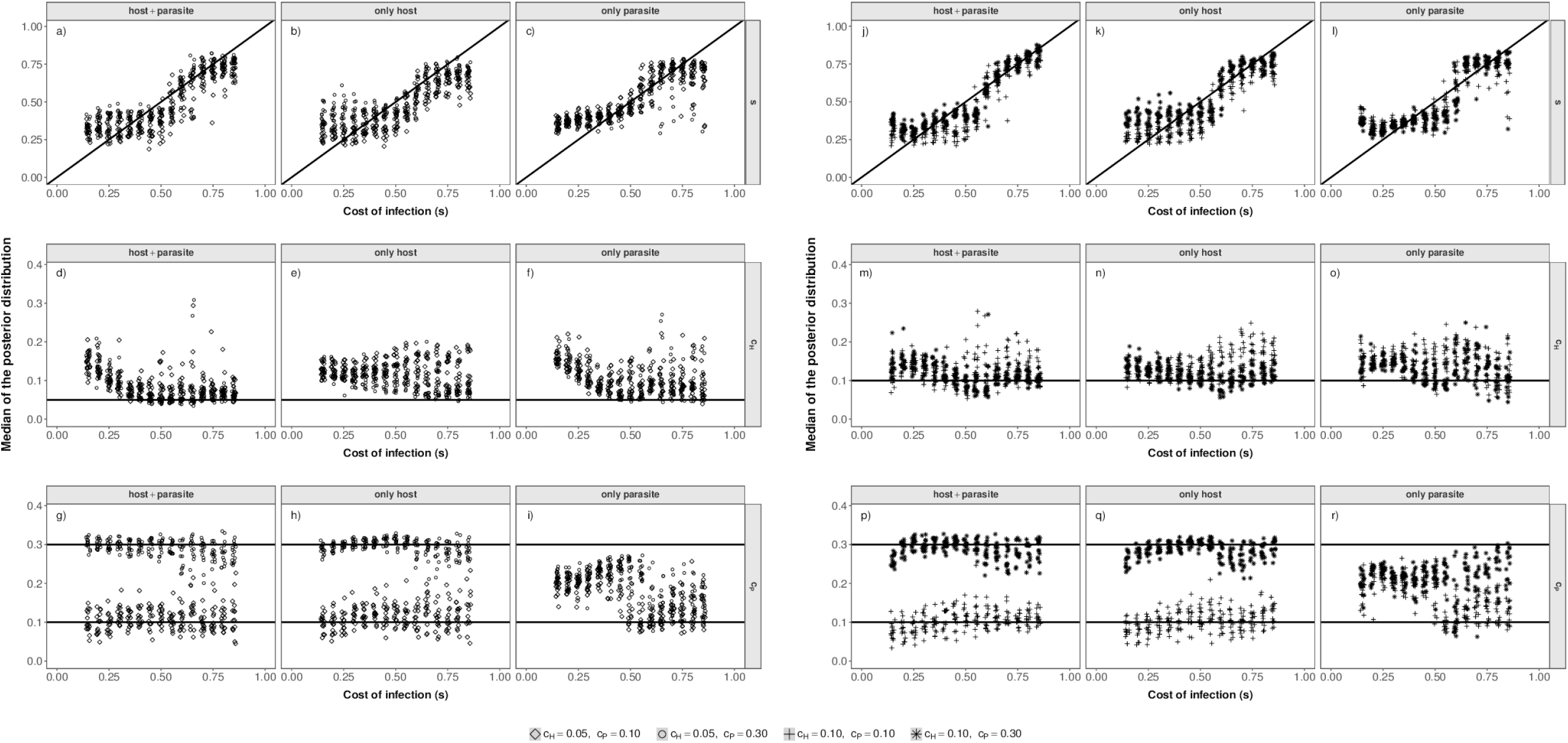
Parameter estimations under scenario 2. Median of the posterior distribution (y-axis) for the cost of infection *s* (top, a-c), cost of resistance (*c*_*H*_) (middle, d-f) and cost of infectivity (*c*_*P*_) (bottom, g-i) when inference is based on host and parasite summary statistics (left), only host summary statistics (middle) or only parasite summary statistics (right). The median of the posterior distribution (after post-rejection adjustment) is plotted for each POD. The true cost of infection for each POD is shown on the x-axis.

## Discussion

In the present study we explicit a link between coevolutionary dynamics (S2 Fig), the resulting genetic signatures (S9 Fig, S3 Fig) and the subsequent amount of information which can be extracted from genetic signatures at the coevolving loci (Fig 3, Fig 6). Our results indicate that under trench-warfare dynamics the allele frequencies at the non-trivial internal equilibrium point affect the strength of genetic signatures at the coevolving loci in both, the host and parasite. Furthermore, pairs of coevolving loci are well discriminated from pairs of neutral loci by ABC model choice (Fig 1, Fig 2, S6 Fig, S12 Fig), while the accuracy decreases for higher costs of infection. We further show as a proof of principle that it is possible to infer the parameters underlying the coevolutionary interaction from polymorphism data at the loci under coevolution if some relevant parameters such as diverse costs (Fig 3) or population sizes (Fig 6) are known. The inference is accurate if polymorphism data from both the host and the parasite are available from at least ten repetitions of the coevolutionary history (S8 Fig, S14 Fig).

### Coevolutionary dynamics and inference

As already shown in [19] there is a continuum of genetic signatures which can arise at the loci under coevolution. This contrasts to the often postulated dichotomy that arms-race dynamics result in strong selective sweep signatures and trench-warfare dynamics in strong balancing selection signatures.

In general, the strength of the selective signatures under trench-warfare dynamics is a result of the internal equilibrium frequencies, the fluctuations around these equilbrium frequencies, the amount of genetic drift in both partners and the proximity of these equilibrium frequencies to the fixation boundaries. When equilibrium frequencies are close to boundaries, alleles can be easily lost by drift and thus, arms-race dynamics take place although trench-warfare dynamics would be predicted based on the deterministic model.

The strong link between equilibrium frequencies under trench-warfare dynamics and resulting genetic signatures can be explained in terms of the underlying approximated structured coalescent tree with two alleles in each species (*RES* and *res* for the host and *INF* and *ninf* for the parasite). This model is analogous to a two-demes model with gene flow [43]. When the frequencies of both allles are fairly similar they have equal contributions to the sample, and the underlying coalescent tree is well balanced. Accordingly, we observe an excess of intermediate frequency variants in the SFS ([11]). As the equilibrium frequencies move away from 0.5, the average sample configuration changes and the coalescent tree becomes less balanced (see S1 Fig). Therefore, the number of SNPs at intermediate frequencies drops and Tajima’s D decreases (S9 Fig). This link can be also observed when we modify our model to more realistic but complex models by either a) extending the model to more than two parasite generations per host generation (**Model B**, S15 Fig, S17 Fig a+b) or b) allowing for allo-infections at rate 1 – *ψ* in the second parasite generation within host generation *g* (**Model C**, S16 Fig, S17 Fig c+d).

There are three sources of stochasticity affecting the polymorphism data at the coevolutionary loci: 1) The effect of genetic drift on the allele frequency trajectory under coevolution, 2) the stochasticity in the coalescent process for a given allele frequency trajectory and 3) the stochasticity in the neutral mutation process on top of the coalescent process. As the first type of stochasticity affects the ‘population’ sizes of the functional alleles in the host (in the parasite) over time, it also has a subsequent effect on the other two sources of stochasticity. Using data from several repetitions allows to better handle and to average out the effect of genetic drift on the variability of the allele frequency path and its subsequent effect on the observed summary statistics. Therefore we use the average of the summary statistics over several repetitions of the same coevolutionary history (*i.e. r* frequency paths) in our ABC. In future, repeated data could be for example obtained from microcosm experiments such as performed by [54,55]). Using data from repeated experiments is one possible attempt to deal with the variability in allele frequency trajectories. The availability of data from several independent populations or time-samples might be possible alternatives. Time samples offer a partially detailed view on changes in allele frequencies and accordingly, can help to better capture the coevolutionary dynamics.

### Accuracy of inference

We first perform a model choice procedure for each scenario to assess the possibility to distinguish pairs of loci which are coevolving from pairs evolving independently from one another (in our case neutrally in each species). We envision that the gene dataset can be divided into two categories of genes in hosts and parasites: pairs of candidate loci possibly under coevolution, and pairs of other randomly selected genes. For example, the candidates can be resistance genes in the host plant [4–6,12–16] and the corresponding predicted effectors in the parasite [17,18]. The second category can be composed of genes involved in processes such as housekeeping, abiotic stress responses or photosynthesis in plants, and housekeeping genes and/or degrading enzymes with non-specific activities in parasites.

It is encouraging that our results show very good accuracy and low False Positive rates. Interestingly, the model choice accuracy is very good for low values of the infection parameter *s*, and thus we are more likely to identify pairs of loci which are coevolving under trench-warfare than under arms race dynamics. We show thus that in contrast to the somehow pessimistic view in [19] based on few statistics, extending the number of summary statistics does help to distinguish neutral from coevolving loci.

Regarding parameter inference, we show that estimations of parameters governing the coevolutionary dynamics is possible if they substantially shift the equilibrium frequencies and/or the dynamics and thus, the resulting genetic signatures. However, equilibrium frequencies can be shifted along the same axis by different parameter combinations. In such circumstances, it is only possible to infer a compound parameter if there is no *a priori* information on any of the parameters available. This identifiability problem is illustrated by the inference results for scenario 2 especially when only parasite polymorphism data are available (Fig 6). Here, both the cost of infection (*s*) and the cost of resistance (*c*_*H*_) are overestimated. If however some parameter values are *a priori* known from experiments such as the cost of resistance in scenario 1, the other parameters (here the cost of infection) can be inferred conditional on this information. Whenever the parameters of interest have different effects on the equilibrium frequencies in the host and parasite, inference of both parameters is possible. This explains why inferences are usually the most accurate when host and parasite statistics are jointly used.

Our approach of jointly using host and parasite information is in line with recent method developments [56–58] which also show the value of analysing hosts and parasite in a joint framework. These mentioned methods can be promising complementary approaches to our ABC in order to identify candidate loci.

### Scope, implications and applications of the presented approach

Based on the genetic signatures found for our two model extensions (S17 Fig) [24], we suggest that our findings are generally valid and are not restricted to the coevolution GFG model used in the main text. We acknowledge that we assume the most simple type of coevolutionary interaction possible. However, understanding possible links between dynamics, signatures and resulting accuracy of inference for this simple scenario is a useful starting point to develop further inference methods where several major loci [7] or quantitative traits [39] are involved. In addition, our approach should be applicable to several pairs of host and parasite coevolving loci as long as the coevolutionary dynamics are driven by few major loci without any epistatic and/or pleiotropic effect. These pairs could for example involve resistance genes from a single host species, each co-evolving independently with effectors from different parasite species (bacteria, fungi, …). If quantitative traits [7,39] are involved into coevolution we expect the signatures to be weaker than in our model (see theory on polygenic selection and polymorphism signatures, [59]).

For many host-parasite models (including the one used here) it has been shown that the equilibrium frequencies in the host are substantially or exclusively affected by fitness penalties applying to the parasite and vice-versa [24,33,34]. Thus generally speaking, the strength of genetic signatures in either species are presumably most indicative about processes affecting the coevolving partner. We therefore speculate, that the balancing selection signatures which have been found at R-genes in *Arabidopsis thaliana* [6,12] [13], *Solanum sp.* [4,5,14], *Phaseolus vulgaris* [60], *Capsella* [61], are indicative about the selective pressure in the coevolving parasite or parasite community. Conversely, the long term maintenance of strains in *Pseudomonas syringae* [62] could reflect fitness costs in *Arabidopsis thaliana*.

A final complication for analysis is the lack of recombination in genomes of microparasites such as viruses or bacteria. Phylogenetic methods exist to study the evolution of these parasites with very short generation time, and can allow to define groups of individuals or populations which could be used in inference methods such as ours or in co-GWAs [56–58,63]. Note also that several methods have been developed to draw inference of the epidemiological parameters based on parasite sequence data (*e.g.* [64]). However, such methods study only short term epidemiological dynamics within few years, ignoring the effect of coevolution and Genotype (host) x Genotype (parasite) interactions. By contrast, our method intends to infer the parameters of long term coevolutionary dynamics driven by GxG interactions.

### Additional demographic changes

An important assumption of our model is the absence of intra-locus recombination at the coevolutionary loci. Nevertheless, recombination does occur along the genomes of the host and the parasite, so that the coevolutionary loci evolve independently from other unlinked loci (for example on different chromosomes).

In such circumstances, it is possible to estimate past population size fluctuations based on whole-genome data of both species. Population size changes in host-parasite coevolution can be either independent of the coevolutionary interaction or arise as an immediate result of coevolutionary interaction, e.g. from epidemiological feedback or any other form of eco-evolutionary feedback. Independently of the particular source, demographic changes always affect all loci in the genome simultaneously. The genomic resolution of the latter type of population size changes has been shown to depend on the amplitude and time-scales of the population size fluctuations [65]. These authors have demonstrated that populations size fluctuations only leave a signature in the genome-wide parasite site frequency spectrum if they happen at a slow enough time scale. Irrespective of whether the demographic changes can be resolved from genome-wide data, the resulting genetic signatures at the coevolving loci will be always the result of underlying allele frequency path which is always confined to a 2d-plane for a bi-allelic locus. Further studies should therefore focus on the specific effect of eco-evolutionary feedback on the variability of the allele frequency path and the resulting effect of the population size changes on mutation supply at the coevolving loci. Doing so will help to refine our understanding how much information can be likely inferred under such circumstances.

### Conclusion

We investigated here a link between coevolutionary dynamics and resulting genetic signatures and quantify the amount of information available in polymorphism data from the coevolving loci. Although, we started from a very simple coevolutionary interaction we show that model-based inference is possible. With growing availability of highly resolved genome data, even of non-model species, it is important to gain a differentiated and deep understanding of the continuum of possible links between coevolutionary dynamics without or with eco-evolutionary feedbacks and their effect on polymorphism data. Such thorough understanding is the basis for devising appropriate sampling schemes, for using optimal combinations of diverse sources of information and for developing model-based refined inference methods. Our results and the suitability of the ABC approach open the door to further develop inference of past coevolutionary history based on genome-wide data of hosts and parasites from natural populations or controlled experiments. Lastly, as the false positive rate to detect genes under coevolution is smaller than 2.5% (*r* = 30) under the model choice procedure, our method can be used as a starting point to identify host and parasite candidate loci for further functional studies.

## Materials and methods

### General outline of the approach

Approximate Bayesian computation (ABC) is an inference method which can be used in situations where likelihood calculations are intractable, as is the case for the coevolutionary models [39]. The principle of ABC methods is to perform a large amount of simulations covering the parameter space for each of several possible models which are expected to reflect the past evolutionary history of the population(s) of concern and thus having given rise to the observed data. These values of the different parameters of each model are drawn from prior distributions based on current knowledge. The observed data and each simulation are summarised by the same set of summary statistics to reduce their dimensionality. In a rejection step the best set of simulations, the simulations with the smallest distance to the summary statistics of the observed data, can be selected. Based on this retained simulations a model choice can be applied to obtain a posterior probability for each competing model. Under the model with the highest posterior probability, an additional regression step can be used to generate the posterior distribution of each parameter. In this paper we do not use real observed sequence data, but study the power of our approach using so-called pseudo-observed datasets.

In more detail the workflow in our paper is as follows:

1. We compare a model of coevolution between a single host and single parasite locus to a neutral model of independently evolving (non-interacting) pairs of host and parasite loci. Under each model, we simulate polymorphism data for *n* = 50 haploid host individuals and *n* = 50 haploid parasite individuals.
2. We simulate *r* replicates of these data corresponding to repeating *r*-times the coevolutionary history. Such repetitions can be obtained in controlled laboratory set-ups using for example microcosm/chemostat experiments with several replicates, or from several independent natural populations of the same host-parasite system with similar environmental conditions.
3. We summarise the obtained SNP data by a set of 17 statistics for each of the *r*–replicates.
4. We calculate the mean for each of the 17 statistics across the *r*-replicates. These average values are used as summary statistics in the ABC. Therefore, one set of *r* replicates defines a given pseudo-observed dataset (POD).
5. We first perform a model choice between the coevolution model and the neutral model based on our PODs. For each POD, we select the 1% closest simulations based on the set of summary statistics. Based on these retained simulations we compute the posterior probability of both models.
6. In a second step, we estimate the posterior distribution of the coevolutionary parameters for the PODs. We apply a post-sampling adjustment (regression) based on the 1% best simulations under the coevolutionary model.

### Simulation of SNP data at the coevolutionary loci

SNP data at the coevolutionary loci are simulated by using a forward-backward approach as outlined in [19].

#### Forward in time coevolution model

We model coevolution between a single haploid host and a single haploid parasite species. The coevolutionary interaction in both species is driven by a single bi-allelic functional site (SNP, indel, …). This functional site is located in the coevolutionary locus which encompasses several other neutral sites. Hosts are either resistant (*RES*) or susceptible (*res*) and parasites are either non-infective (*ninf*) or infective (*INF*). Thus, the model follows a gene-for-gene interaction with the following infection matrix:

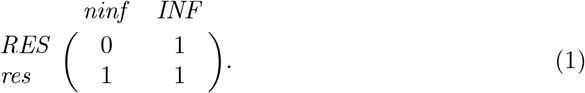

A 1-entry in the infection matrix indicates that the parasite is able to infect the host and a 0-entry indicates that the host is fully resistant towards the parasite. We denote the frequency of resistant hosts (susceptible hosts) by *R* (*r*) and the frequency of infective parasite (non-infective parasites) by *a* (*A*). The coevolution model is based on the polycyclic auto-infection model in [24]. This population genetics model (*sensu* [40]) assumes host and parasite population sizes to be constant regardless of the disease prevalence and is based on non-overlapping host and parasite generations. As such it is probably most suited to describe plant-parasite or invertebrate-parasite systems. Polycyclic diseases are characterized by more than one infection cycle per season. For simplicity, the model is based on *T* = 2 infection cycles per discrete host generation *g* each caused by a single discrete parasite generation *t* (*t* ∈ {1, 2}). An auto-infection refers to an infection where a parasite re-infects the host individual on which it was produced. Therefore, resistant (*R*_*g*_) and susceptible hosts (*r*_*g*_) which are infected by infective parasites (*a*_*g*,1_) in the first infection cycle (*t* = 1) stay infected by infective parasites in the second infection cycle (*t* = 2). This causes a fitness reduction *s*_1_ = *s* (cost of infection). The same applies to susceptible host (*r*_*g*_) infected by non-infective parasites (*A*_*g*,1_) in the first infection cycle (*t* = 1). Resistant host which are attacked by non-infective parasites in the first infection cycle (*t* = 1) resist infection. In the second infection cycle (*t* = 2), this fraction of resistant hosts (*R*_*g*_ · *A*_*g*,1_) either receives a non-infective parasite (*A*_*g*,2_) resulting in no fitness loss or an infective parasite (*a*_*g*,2_) resulting in a reduced cost of infection *s*_2_ = *s*/2. Host resistance comes at cost *c*_*H*_ (cost of resistance) and infectivity in the parasite comes at cost *c*_*P*_ (cost of infectivity).

The allele frequencies of resistant hosts (*R*_*g*_), susceptible hosts (*r*_*g*_), non-infective parasites (*A*_*g*,*t*_) and infective parasites (*a*_*g*,*t*_) are given by the following recursive equations:

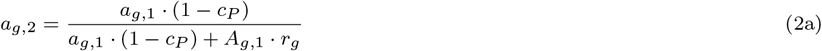

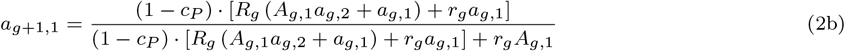

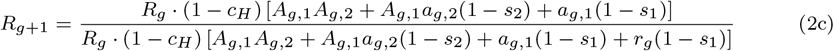

with *A*_*g*,*t*_ = 1 − *a*_*g*,*t*_ and *r*_*g*_ = 1 – *R*_*g*_. The equilibrium frequencies *â*, 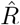 [24] at the internal, non-trivial equilibrium point are approximately given by:

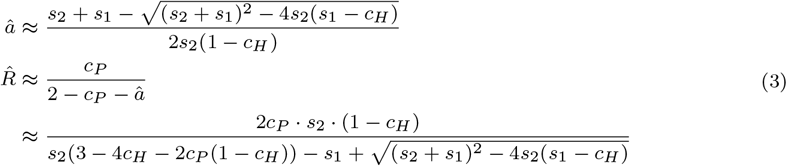

In the forward part, we obtain the frequencies of the different alleles at the beginning of each discrete host generation *g* in three steps:

1. Using the discrete-time gene-for-gene coevolution model from Eq (2), we compute the expected allele frequencies in the next generation (under the infinite population size assumption).
2. Genetic drift is incorporated by performing a binomial sampling based on the frequency of the *RES*-allele (*INF*-allele) after selection (Eq (2)) and the finite and fixed haploid host population size *N*_*H*_ (parasite population size *N*_*P*_) as in [19] (see [41]).
3. Recurrent allele mutations take place and change genotypes from *RES* to *res* at rate *µ*_*Rtor*_ or *res* to *RES* at rate *µ*_*rtoR*_ in the host and from *ninf* to *INF* at rate *µ*_*ntoI*_ and from *INF* to *ninf* at rate *µ*_*Iton*_ in the parasite. Henceforward, such mutations are referred to as functional mutations. In the reminder of this manuscript we set all functional mutation rates to *µ*_*Rtor*_ = *µ*_*ntoI*_ = *µ*_*rtoR*_ = *µ*_*Iton*_ = 10^−5^ (for a discussion on these values see [19,41]).

Repeating this procedure for *g*_*max*_ host generations, we obtain the so called frequency path, which summarizes the allele frequencies at both loci forward in time.

#### Backward in time coalescent

To obtain polymorphism data at the coevolutionary loci we combine the obtained frequency paths which include genetic drift and recurrent mutations with coalescent simulations separately for the host and the parasite. The host and parasite frequency paths are used separately as input for a modified version of msms [19,42], after scaling time appropriately in units of the respective population sizes (for more information see S1 File). Based on the allele frequency in a species at present, a coalescent tree is build backward in time using msms. A sample of size *n*_*H*_ (*n*_*P*_) is drawn at random from the host (parasite) population consisting of *RES* and *res*-alleles (*ninf* and *INF*-alleles) [19]. The tree shape and length are conditioned on the changes in allele frequencies, including fixation or loss [19]. To clarify the forward - backward correspondence, let us describe the case of recurrent selective sweeps in the parasite population. In a monomorphic parasite population of allele *INF*, a functional mutation with rate *µ*_*Iton*_ can reintroduce forward in time a mutant *ninf*. This allele reaches fixation and the population is then monomorphic for allele *ninf*. Backward in time, this is equivalent, in msms, to the decrease of the *ninf* allele population size until only one last individual exhibits this allele. This last *ninf* coalescent lineage then migrates to the population of allele *INF*. The forward frequency path and the backward msms simulations are thus coupled for the re-introduction of new alleles due to functional mutations in analogy to gene flow in a structured coalescent with two demes [43].

The forward in time coevolution model is run for *g*_max_ = max(3*N*_*H*_, 3*N*_*P*_) generations assuming small initial frequency of *RES* (*R*_0_ = 0.2) and *INF* (*a*_0_ = 0.2) alleles. The length of simulation time was previously found to be sufficient to observe signatures of selective sweeps and balancing selection in host or parasite [19]. In msms, the backward simulations conditioned on the frequency paths are run for the same amount of time. If after *g* generations, several coalescent lineages remain and/or the most recent common ancestor of both functional alleles has not been reached, a neutral Kingman coalescent process is built until a common ancestor of all remaining lineages is found. Note that that in this last temporal phase of the simulation, *i.e.* older than *g* generations in the past, the functional alleles in hosts (*RES* and *res*) and in parasites (*INF* and *ninf*) have the same fitness (and are exchangeable within species). We therefore simulate a coevolution history of *g* generations.

We set the sample size to *n*_*H*_ = 50 for the host (*n*_*P*_ = 50 for the parasite) which are adequate to capture balancing selection if one of the allele occurs in low frequency at the present time of sampling [19]. For both species we assume realistically a locus of length 2500 bp without recombination and a per site neutral mutation rate of 10^−7^. Accordingly, the neutral population mutation rate is *θ*_*H*_ = 2 · *N*_*H*_ · 2500 · 10^−7^ for the host (*θ*_*P*_ = 2 · *N*_*P*_ · 2500 · 10^−7^ for the parasite) defining the number of mutations found on the host and parasite coalescent trees (and in the polymorphism data).

#### Calculating statistics for the SNP-data

For each msms-output we calculate eight statistics for each species which are based on the site frequency spectrum (SFS) of the respective coevolving locus (Tab. 1). We only use statistics based on the unfolded site frequency spectrum (SFS), as it can be hard to obtain unbiased haplotype statistics depending on the sequence method. In addition to these 16 statistics we calculate the (**P**airwise **M**anhattan **D**istance) which is based on comparing the host and parasite site frequency spectra (see S2 File).

**Table 1.** SNP statistics calculated.

| Name | reference |
| --- | --- |
| number of segregating sites $S$ | [44] |
| $\theta_W$ | [44] |
| nucleotide diversity $\pi$ | [45] |
| Tajimas' D | [46] |
| Fu and Li's D | [47] |
| Fu and Li's F | [47] |
| $\theta_H$ | [48] |
| Hprime | [49] |
| PMD |  |

#### Additional coevolutionary models tested

Additionally, we study two extensions, **B** and **C** of the model from Eq (2) (Model **A**), in order to check for the generality of our results. In model **B**, we extend the described model to include more than two parasite (*T* > 2) generations per host generation *g* (see S1 File). In model **C**, we keep *T* = 2 but allow for allo-infection to take place at rate (1 − *ψ*) in the second parasite generation (*t* = 2) within host generation *g* (see S1 File). Based on the equations (S1 File), we generate forward in time simulations with genetic drift and functional mutations (as described above) and the expected coevolutionary signatures at the coevolving loci. We study how the values of the different statistics obtained under these two more realistic but complex models differ from those of the main model from Eq (2).

### ABC inference

In the following section, we lay out the two scenarios to be investigated, the simulations for obtaining the PODs, and the prior distributions for the coevolutionary and neutral models. Finally, the ABC model choice and parameter estimation procedures are described.

#### Inference scenarios

We focus on two scenarios. In scenario 1, we aim to infer the cost of infection (*s*), the host population size (*N*_*H*_) and the parasite population size (*N*_*P*_). Therefore, the cost of resistance (*c*_*H*_) and the cost of infectivity (*c*_*P*_) are assumed to be known. In scenario 2 the goal is to infer the cost of infection (*s*), the cost of resistance (*c*_*H*_) and the cost of infectivity (*c*_*P*_), assuming that the host (*N*_*H*_) and parasite (*N*_*P*_) population sizes are known.

#### Generating pseudo-observed data sets

Each pseudo-observed datasets (PODs) is composed of *r* = 30 repetitions of the coevolutionary history under a particular combination of parameters (*s*, *c*_*P*_, *c*_*H*_) while fixing the haploid population sizes to *N*_*H*_ = *N*_*P*_ = 10,000 and the population mutation rates to *θ*_*H*_ = *θ*_*P*_ = 5.

For scenario 1, we simulate PODs for values of cost of infection (*s*) ranging from *s* = 0.15 to *s* = 0.85 (in steps of size 0.05) while fixing the cost of resistance to *c*_*H*_ = 0.05 and the cost of infectivity to *c*_*P*_ = 0.1. For each value of *s*, 30 independent PODs are simulated.

For scenario 2, we generate PODs for the 60 possible combinations of *c*_*H*_ ∈ {0.05, 0.1}, *c*_*P*_ ∈ {0.1, 0.3} and *s* from 0.15 to 0.85 (in steps of size 0.05). For each of these combinations, 15 PODs are generated.

#### ABC sampling: priors of the coevolutionary model

For both scenarios, between 95,000 and 100,000 datasets are generated from the coevolutionary model based on the following priors (defined with the ABCsampler from ABCtoolbox, Version 1.0, [50]).

In scenario 1, defined with *c*_*H*_ = 0.05 and *c*_*P*_ = 0.1, the cost of infection is drawn from a uniform prior such that 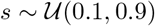, and the host and parasite population sizes are drawn for log uniform distributions such that 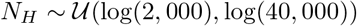 and 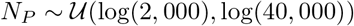. The population mutation rates are calculated as *θ*_*H*_ = 2*N*_*H*_ · 25000 · 10^−7^ and *θ*_*P*_ = 2*N*_*P*_ · 25000 · 10^−7^ (see Tab. 2).

**Table 2.** Settings ABC scenario 1. Settings for the ABC simulations under scenario 1.

|  | Coevolution model | Neutral model |
| --- | --- | --- |
| $N_H$ | $\sim \mathcal{U}(\log(2000), \log(40000))$ | $\sim \mathcal{U}(\log(2000), \log(40000))$ |
| $N_P$ | $\sim \mathcal{U}(\log(2000), \log(40000))$ | $\sim \mathcal{U}(\log(2000), \log(40000))$ |
| $\theta_H$ | $2 \cdot N_H \cdot 2500 \cdot 10^{-7}$ | $2 \cdot N_H \cdot 2500 \cdot 10^{-7}$ |
| $\theta_P$ | $2 \cdot N_P \cdot 2500 \cdot 10^{-7}$ | $2 \cdot N_P \cdot 2500 \cdot 10^{-7}$ |
| $n_H$ | 50 | 50 |
| $n_P$ | 50 | 50 |
| $s$ | $\sim \mathcal{U}(0.1, 0.9)$ | — |
| $c_P$ | 0.10 | — |
| $c_H$ | 0.05 | — |

In scenario 2, defined by *N*_*H*_ = *N*_*P*_ = 10,000 and *θ*_*H*_ = *θ*_*P*_ = 5, the cost of infection is drawn from a uniform distribution such that 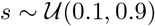, and the cost of resistance and infectivity from uniform distributions such that 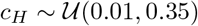 and 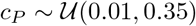 (see Tab. 3).

**Table 3.** Settings ABC scenario 2. Settings for the ABC simulations under scenario 2.

|  | Coevolution model | Neutral model |
| --- | --- | --- |
| $N_H$ | 10,000 | 10,000 |
| $N_P$ | 10,000 | 10,000 |
| $\theta_H$ | 5 | 5 |
| $\theta_P$ | 5 | 5 |
| $n_H$ | 50 | 50 |
| $n_P$ | 50 | 50 |
| $s$ | $\sim \mathcal{U}(0.10, 0.90)$ | — |
| $c_P$ | $\sim \mathcal{U}(0.01, 0.35)$ | — |
| $c_H$ | $\sim \mathcal{U}(0.01, 0.35)$ | — |

#### ABC sampling: priors of the neutral model

As for the coevolution model, we obtain between 95,000 and 100,000 data sets for a corresponding neutral model for each scenario. This neutral simulations are generated by coalescent simulations with msms [42] for a non-recombining host and parasite locus with the same length (2500bp) as in the coevolutionary model. To mimic data obtained from the same repeated evolutionary history, we generate *r* = 30 repetitions of the neutral coalescent process. For each replicate we calculate the same 17 statistics as under the coevolution model which are defined in Tab. 1. The summary values used in the ABC consist of the average over the *r* replicates for each statistic.

Under scenario 1, the neutral simulations are based on priors for the host and parasite population sizes drawn from log uniform distributions 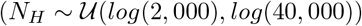 and 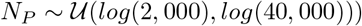. The population mutation rates are calculated as *θ*_*H*_ = 2*N*_*H*_ · 25000 · 10^−7^ and *θ*_*P*_ = 2*N*_*P*_ · 25000 · 10^−7^.

Under scenario 2, we simulate datasets for constant host and parasite population sizes (*N*_*H*_ = *N*_*P*_ = 10,000) and thus the population mutation rates are *θ*_*H*_ = *θ*_*P*_ = 5.

#### ABC model choice

The ABC model choice procedure is used to test whether a pair of coevolving loci can be discriminated from pairs of neutral loci based on our set of summary statistics and within the range of priors for our outlined scenarios. To find genes under coevolution, we wish to access the False Positive (FPR) and the False Negative (FNR) rate. These rates are also referred to as the confusion matrix in the ABC literature. Under the hypothesis that two genes (one from the host and one from the parasite) are coevolving, the FPR is the percentage of pairs of truly neutral loci which have a higher posterior probability in support of the coevolution model rather than the neutral model. Thus, these loci would be incorrectly identified as coevolving although in fact they are not. On the other hand, the FNR is defined as the percentage of truly coevolving pairs of loci which have a higher posterior probability in support of the neutral model (rather than the coevolving model). These loci would be considered as neutral although they are in fact coevolving. To access the FPR and FNR, we first perform a leave-one-out cross-validation running the function cv4postr of the abc r-package (version 2.1, [51]) for each scenario 1 and 2. The cross-validation is based on the rejection algorithm as follows. Under a given scenario (1 or 2), a dataset called validation simulation, is chosen at random from one of the two models (coevolution or neutral). All summary statistics are standardised by their median absolute deviation for all simulations. Based on these normalised summary statistics the Euclidean distance between the summary statistics of the validation simulation and all other simulations from both models is calculated. The one percent of the simulations with the smallest Euclidean distance to the validation simulation are retained and all other simulations are rejected [51]. Based on these retained simulations, the posterior probability for each of the two models is calculated for this given validation simulation. This procedure is repeated for 500 validation simulations for each model within each scenario. The FDR and FNR are thus computed for each scenario. Model choice was also performed for each of the PODs to investigate the effect of specific coevolutionary parameters on the accuracy of model choice. For each scenario we used the same settings and simulations for the coevolution model and the neutral model as for the cross-validation. For each POD we retain the 1% best simulations and report the posterior probability for the coevolution model.

#### ABC parameter estimation

The inference of the coevolution model parameters is obtained using the ABCestimator within the ABCtoolbox (Version 1.0, [50]). We retain the 1,000 simulations with the smallest Euclidean distance (without summary statistics normalisation) to the respective POD (rejection step). The standard ABCestimator applies a Gaussian kernel smoothing for each parameter (width of Dirac peak set to 0.01) followed by a post sampling adjustment via a general linear model [50,52]. We report the median of the posterior marginal density distribution for each parameter. For each POD we perform the parameter estimation based on a) host and parasite summary statistics, b) host summary statistics only and c) parasite summary statistics only.

## Acknowledgments

We thank Amandine Cornille and Cas Retel for helpful comments on early versions of the manuscript and Lukas Heinrich for performing preliminary analyses.

## Supporting information

**S1 Fig.**
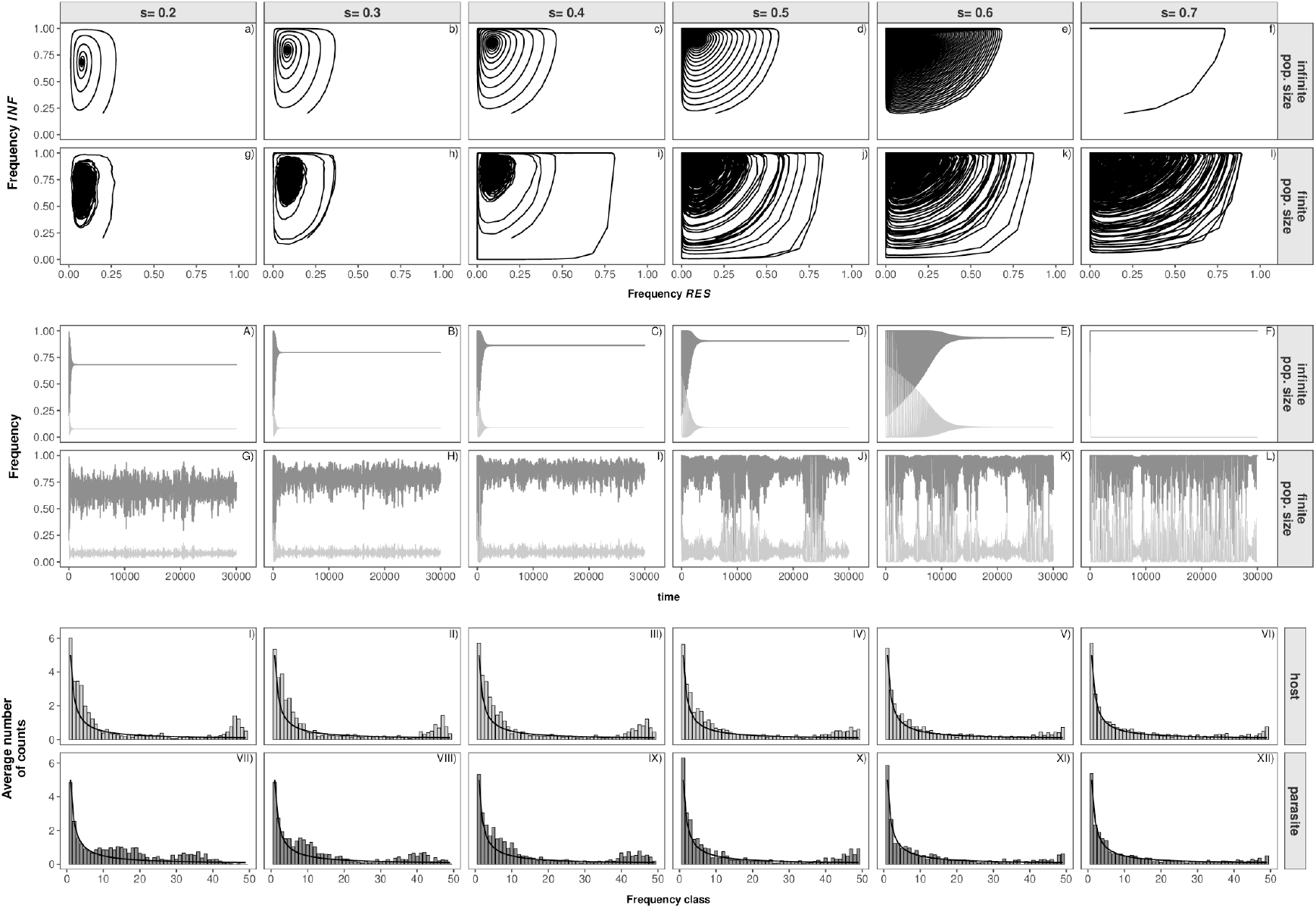
Coevolution dynamics in infinite population size, finite population size and site frequency spectra for Model A. Influence of the cost of infection (*s*) on the coevolutionary dynamics and genetic signatures in **Model A**. The subfigures show the allele frequency trajectory in infinite population size (a-f, A-F), one exemplary allele frequency path in finite population size which takes genetic drift and functional mutations into account (d-f, D-F), the average unfolded host site frequency spectrum of *r* = 200 repetitions (I-VI) and the average unfolded parasite site frequency spectrum of *r* = 200 repetitions (VII-XII). In subfigures a-l each dot represents the frequency of resistant (*RES*) hosts (x-axis) and infective (*INF*) parasites (y-axis) at the beginning of a single host generation *g*. The same information is displayed in a slightly different way in subfigures A-L. Here, the frequencies of resistant (*RES*) hosts (light grey) and infective (*INF*) parasites (dark grey) (y-axis) are plotted over time (x-axis). Costs are fixed to *c*_*H*_ = 0.05, *c*_*P*_ = 0.1. The results in finite population size are plotted for *N*_*H*_ = *N*_*P*_ = 10,000, *µ*_*Rtor*_ = *µ*_*ntoI*_ = *µ*_*rtoR*_ = *µ*_*Iton*_ = 10^−5^. The site frequency spectra are shown for *θ*_*P*_ = *θ*_*H*_ = 5 and *n*_*H*_ = *n*_*P*_ = 50.

**S2 Fig.**
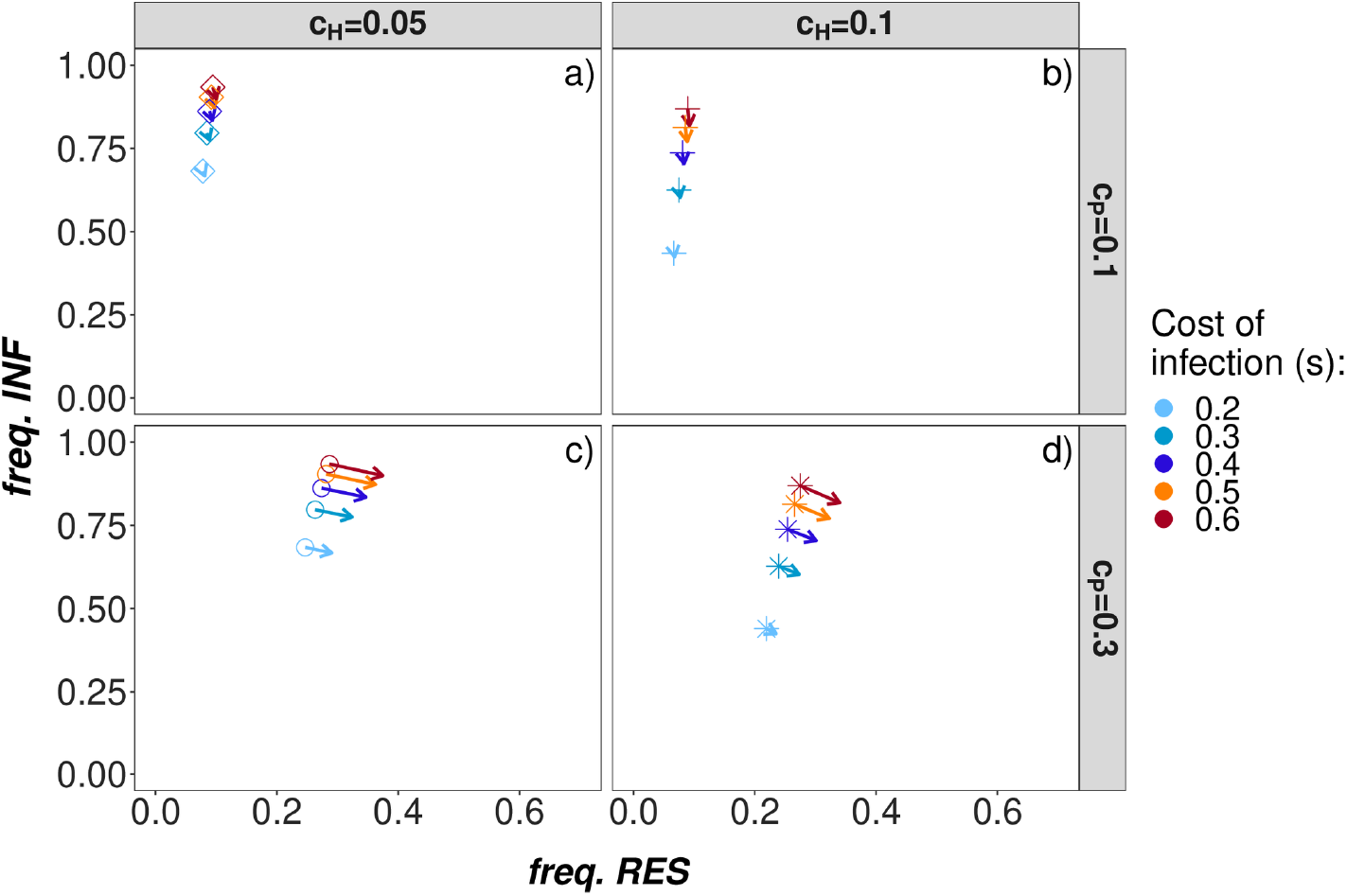
Deterministic equilibrium frequencies model A. Deterministic equilibrium frequencies for model A for different combinations of cost of resistance *c*_*H*_ = (0.05, 0.1) (columns), cost of infectivity *c*_*P*_ = (0.1, 0. 3) (rows) and cost of infection *s* = (0.2, 0.3, 0.4, 0.5, 0.6, 0.7, 0.8) (color of the squares). Only parameter combinations with trench-warfare dynamics are shown. Centres of the dots represent the stable equilbrium frequencies obtained by simulating numerically the recursion equations Eq (2) for 30,000 generations starting with an initial frequency of *R*_0_ = 0.2 resistant hosts and *a*_0_ = 0.2 infective parasites. Heads of the arrows represent the equilibrium frequencies based on Eq (3) which slightly differ from the numerical computations due to analytical approximations.

**S3 Fig.**
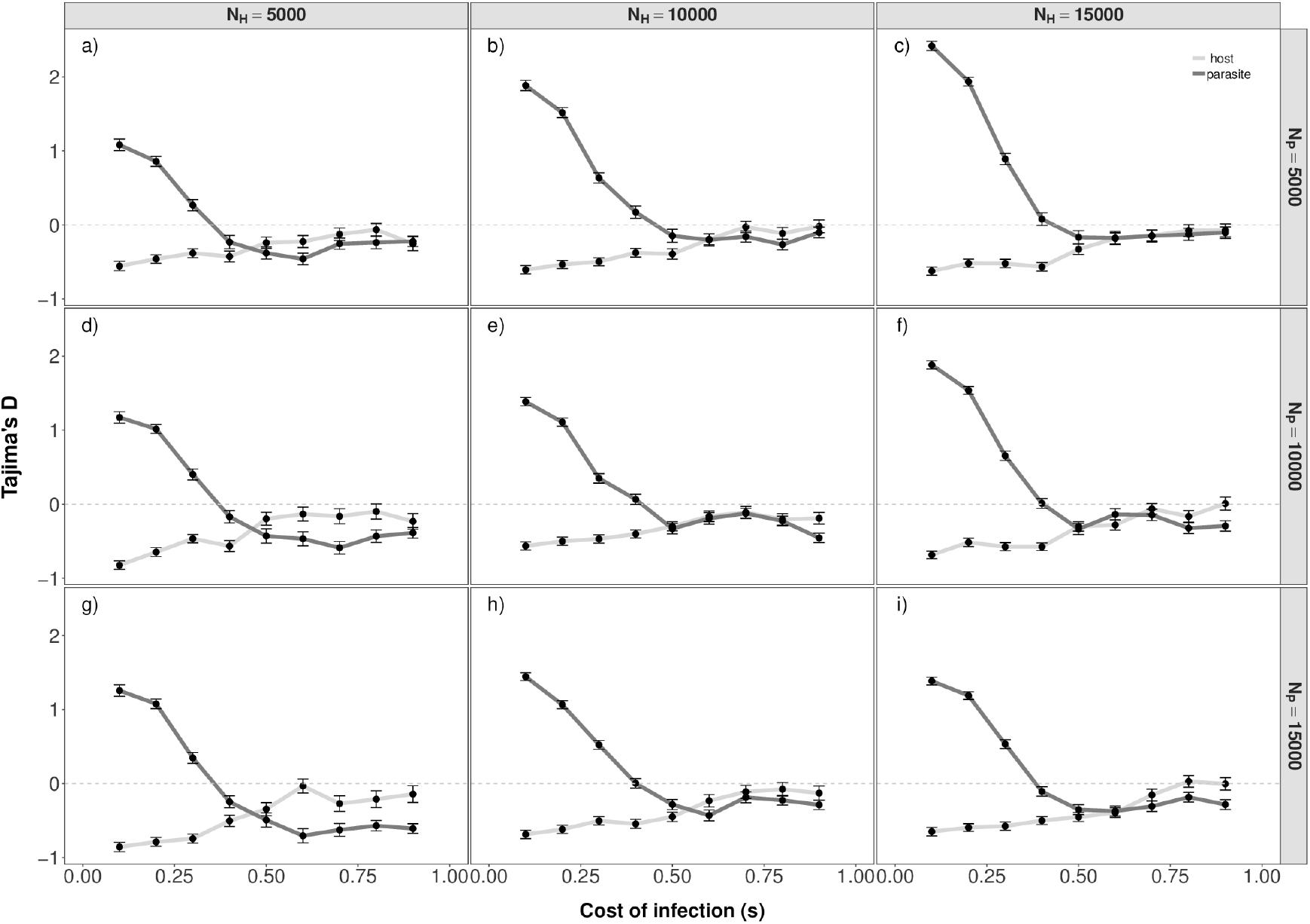
Tajima’s D model A varying popsizes. Tajima’s D (y-axis) for **Model A** for various cost of infection *s* (x-axis) and different combinations of *N*_*P*_ (*N*_*P*_ = 5,000 top, *N*_*P*_ = 10,000 middle, *N*_*P*_ = 15,000 bottom) and *N*_*H*_ (*N*_*H*_ = 5,000 left, *N*_*H*_ = 10,000 middle, *N*_*H*_ = 15,000 right). The mean and standard error of Tajima’s D of the parasite population (dark grey) and of the host population (light grey) are plotted for *r* = 200 repetitions. Note that subfigure *e* corresponds to S9 Fig a. The other parameters are fixed to: *c*_*H*_ = 0.05, *c*_*P*_ = 0.1, *θ*_*H*_ = *N_H_ /*2000, *θ*_*P*_ = *N_P_ /*2000, *n*_*H*_ = *n*_*P*_ = 50, *µ*_*Rtor*_ = *µ*_*rtoR*_ = *µ*_*ntoI*_ = *µ*_*Iton*_ = 10^−5^.

**S4 Fig.**
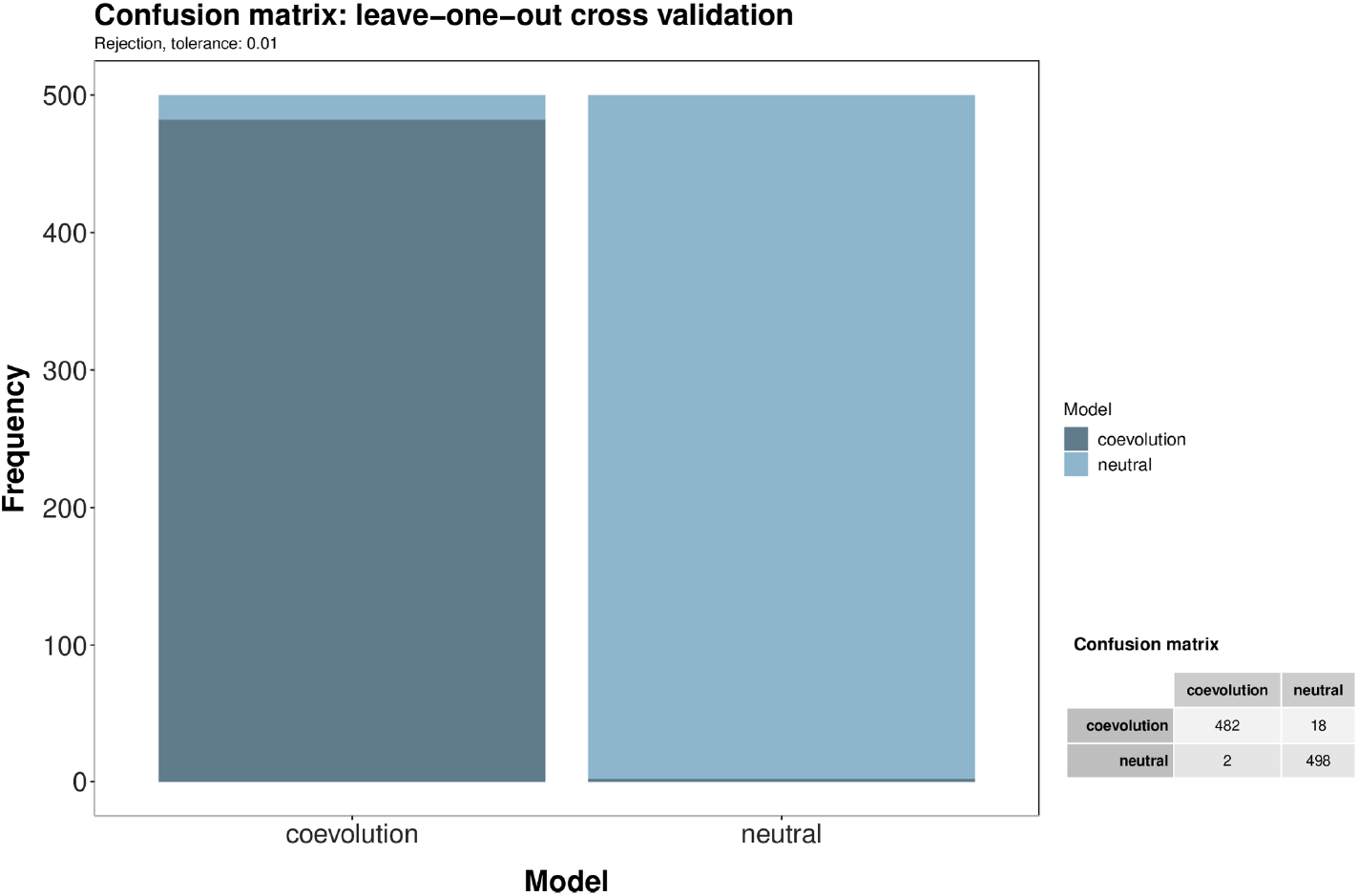
Cross-validation model choice scenario 1 for r = 30 repetitions. Leave-on-out cross-validation result for distinguishing the coevolution model with unknown costs of infection (*s*), host population size (*N*_*H*_) and parasite population size (*N*_*P*_) from a neutral model with a unknown host and parasite population sizes. Cross-validation results are shown for *r* = 30 and are based on 500 randomly chosen ABC-simulations for each model.

**S5 Fig.**
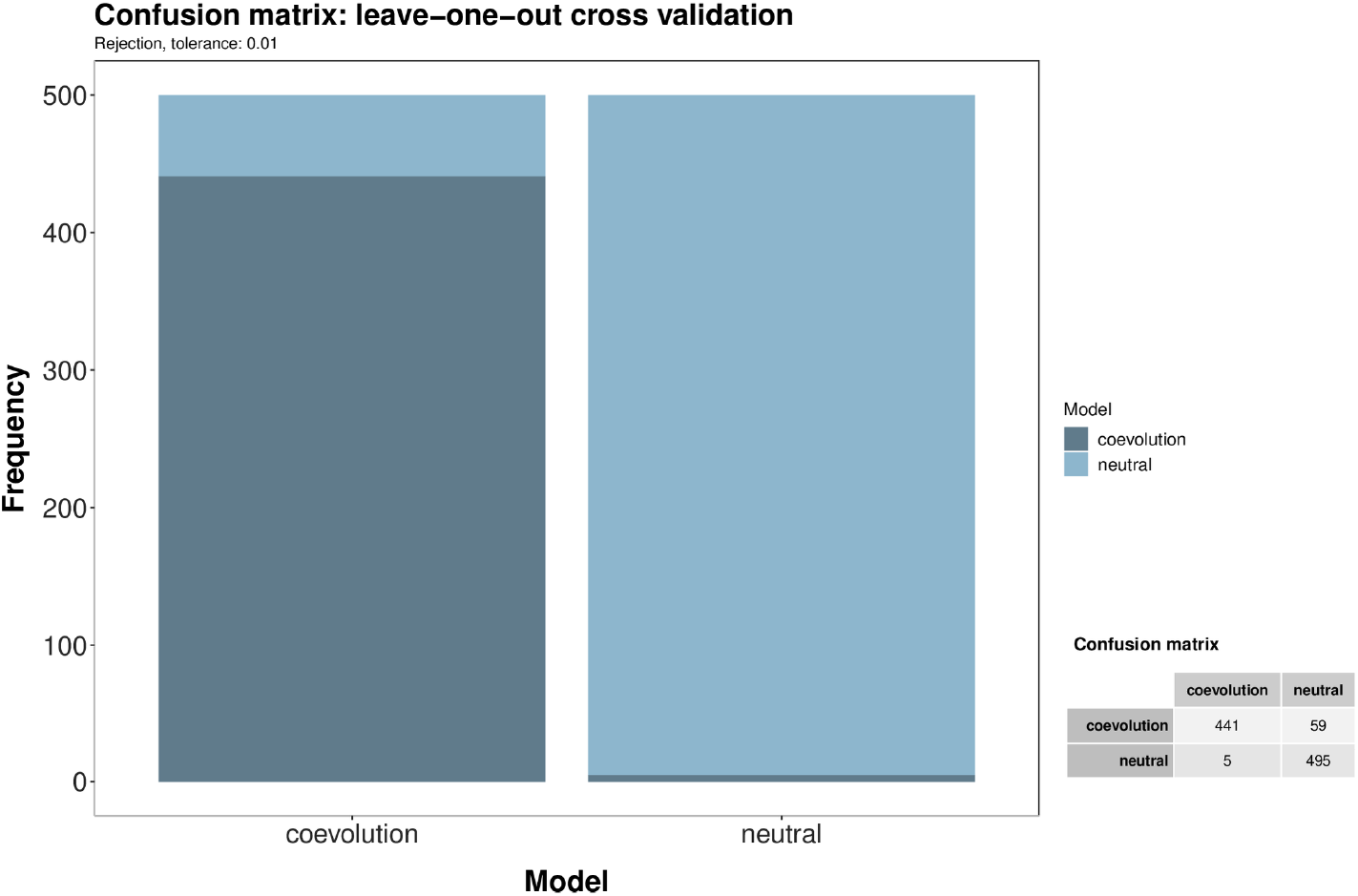
Cross-validation model choice scenario 1 for r = 10 repetitions. Leave-on-out cross-validation result for distinguishing the coevolution model with unknown costs of infection (*s*), host population size (*N*_*H*_) and parasite population size (*N*_*P*_) from a neutral model with unknown host and parasite population sizes. Cross-validation results are shown for *r* = 10 and are based on 500 randomly chosen ABC-simulations for each model.

**S6 Fig.**
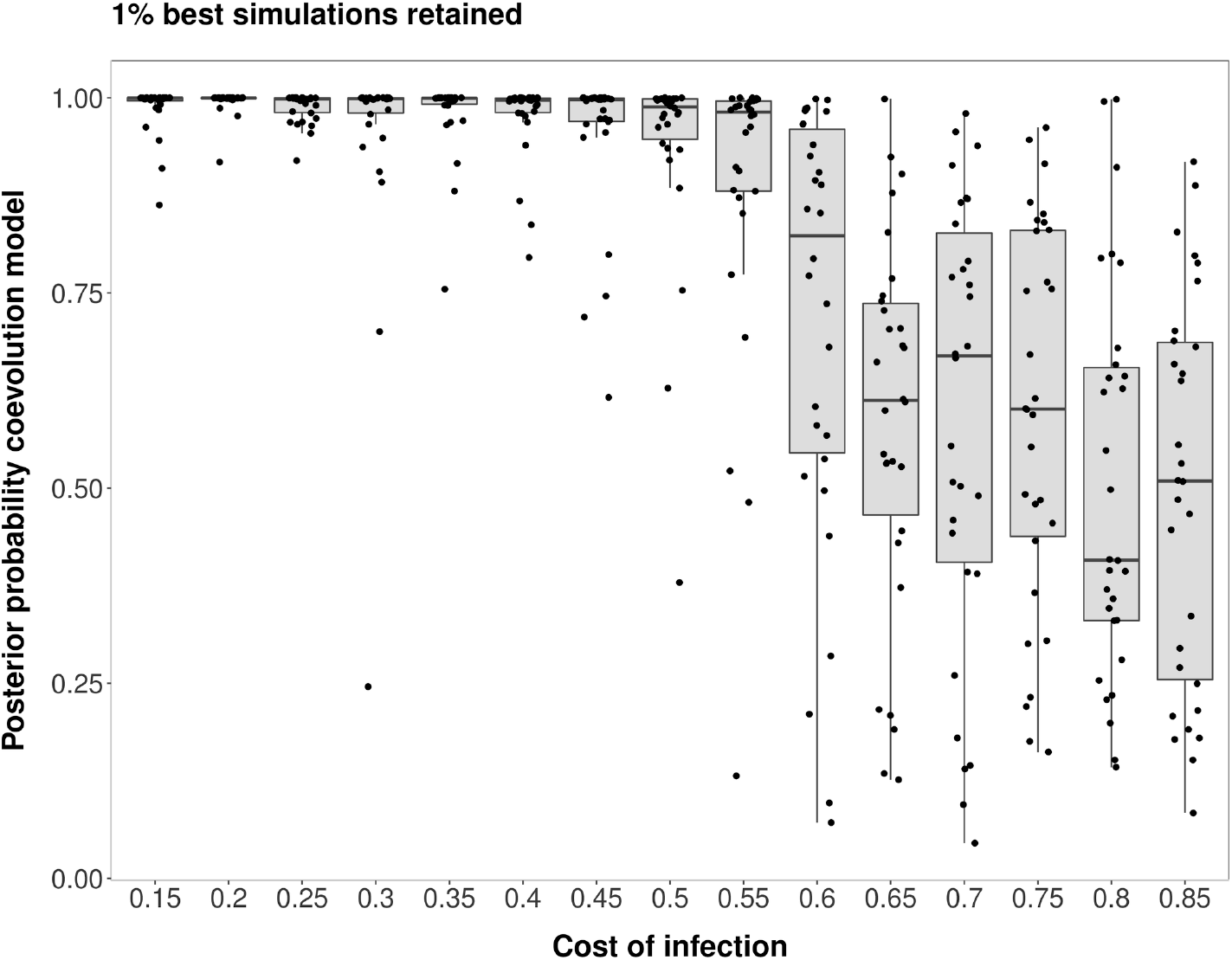
Model choice results for PODs from scenario 1 for r = 10 repetitions. Model choice results for scenario 1 for *r* = 10. Model choice has been run to distinguish a coevolution model with unknown costs of infection (*s*), host population size (*N*_*H*_) and parasite population size (*N*_*P*_) from a neutral model with unknown host and parasite population sizes. Model choice is shown for *r* = 30 repetitions and based on the 1% simulations having the closest summary statistics to those of the PODs. The posterior probability in support of the coevolution model (y-axis) is shown for PODs with different costs of infection (*s*) (30 PODs for each *s*). Results for single PODs are shown as dots. Note that for these points we added some jitter to the x-values in order to increase the readability of the plots.

**S7 Fig.**
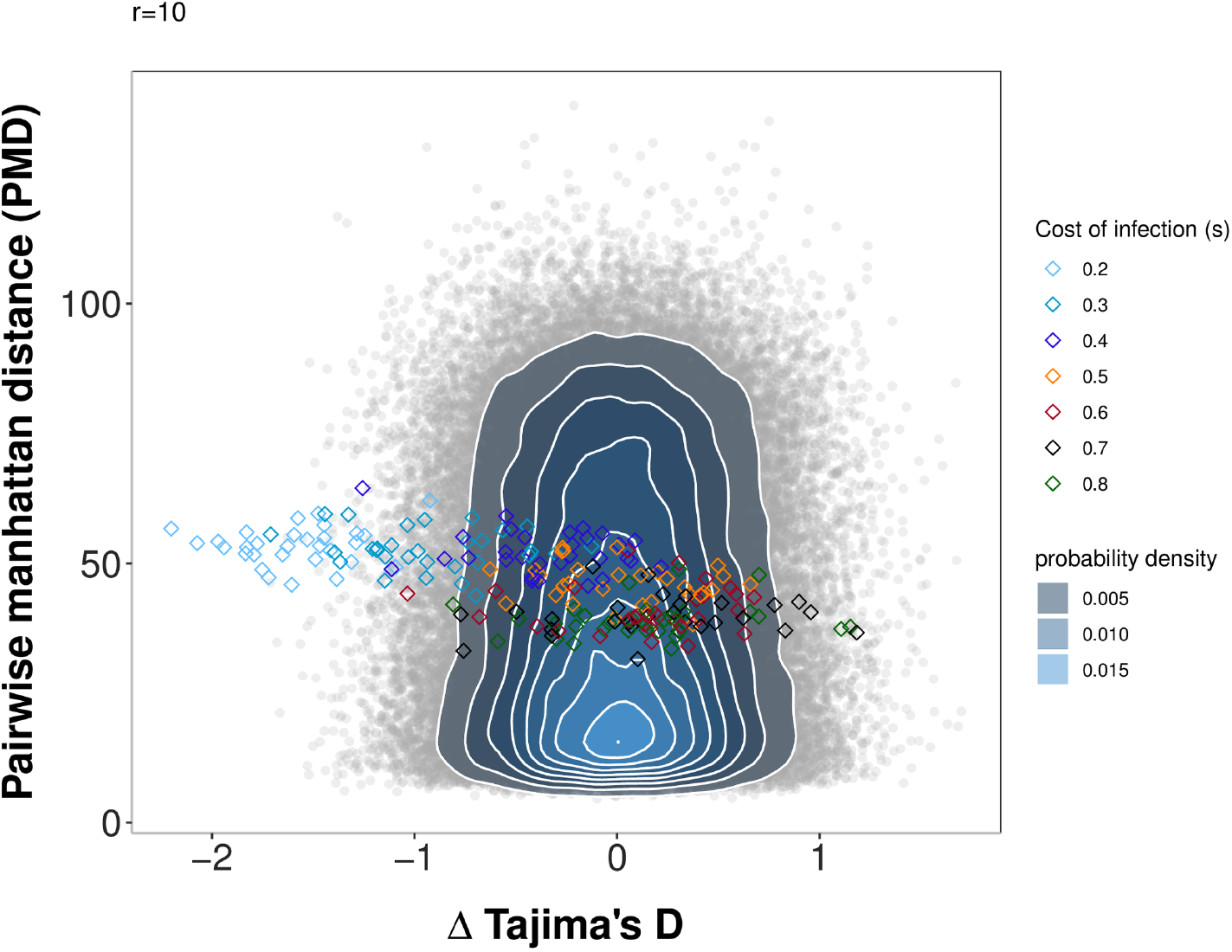
Pairwise Manhattan distance and Δ Tajima’s D (host-parasite) for the PODs under scenario 1 compared to simulations under a neutral model for r = 10. Pairwise Manhattan distance (x-axis) and the difference between Tajima’s D of the host and of the parasite (y-axis) for the PODs used for inference in Scenario 1 and the 100,000 neutral simulations run for this scenario. Under the neutral model, host and parasite population sizes vary. Simulations under the neutral model are shown as grey open circles, and a bivariate normal kernel estimation has been applied to obtain a probability density of the summary statistic combinations. The PODs for scenario 1 are shown as diamonds and are coloured coded based on the true cost of infection (*s*).

**S8 Fig.**
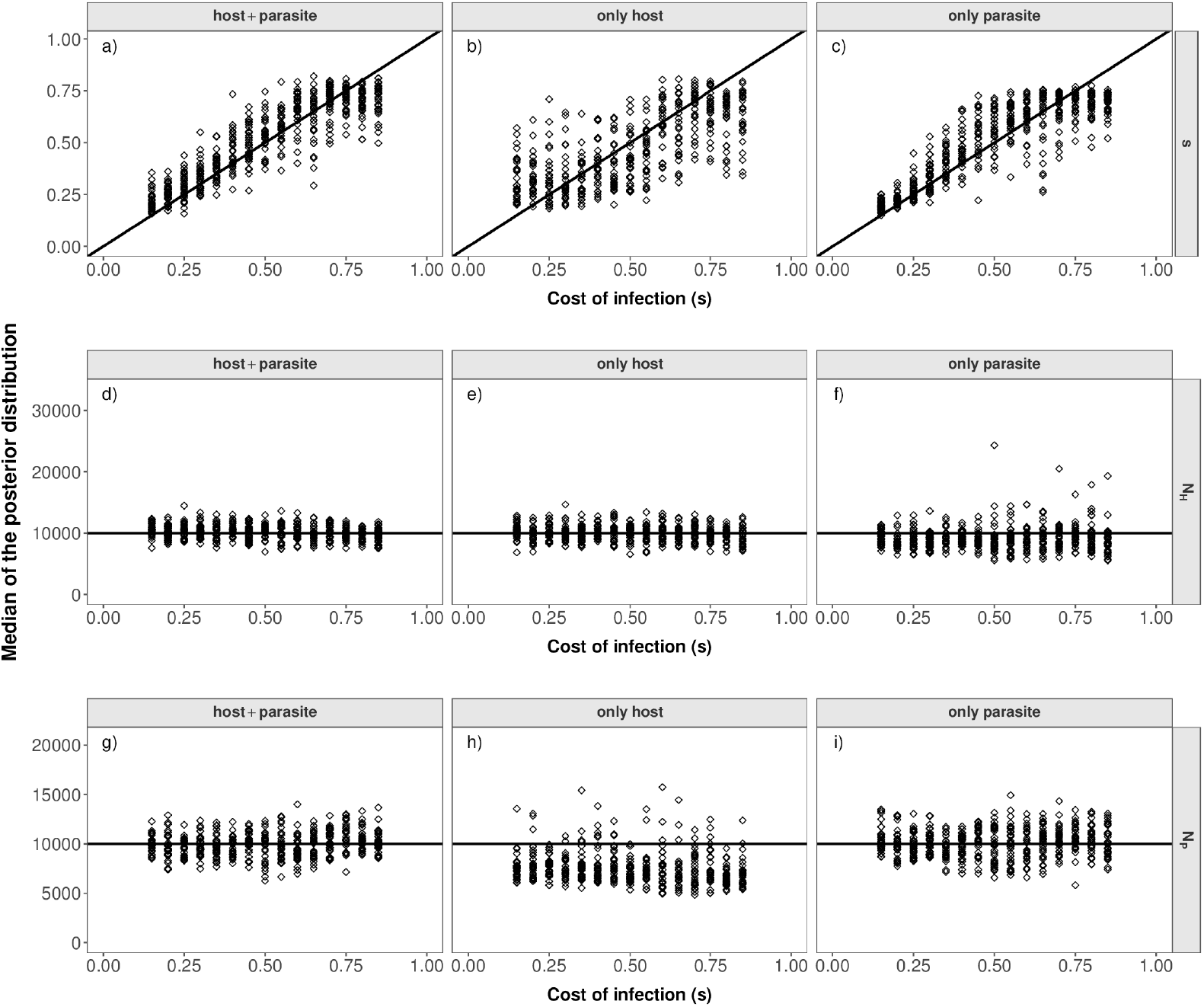
Inference results Scenario 1 for r = 10. Median of the posterior distribution (y-axis) for the cost of infection *s* (top, a-c), host population size (*N*_*H*_) (middle, d-f) and parasite population size (*N*_*P*_) (bottom, g-i) when inference is based on host and parasite summary statistics (left), only host summary statistics (middle) or only parasite summary statistics (right) for scenario 1. The median of the posterior distribution (after post-rejection adjustment) is plotted for each POD in scenario 1. The true cost of infection for each POD is shown on the x-axis.

**S9 Fig.**
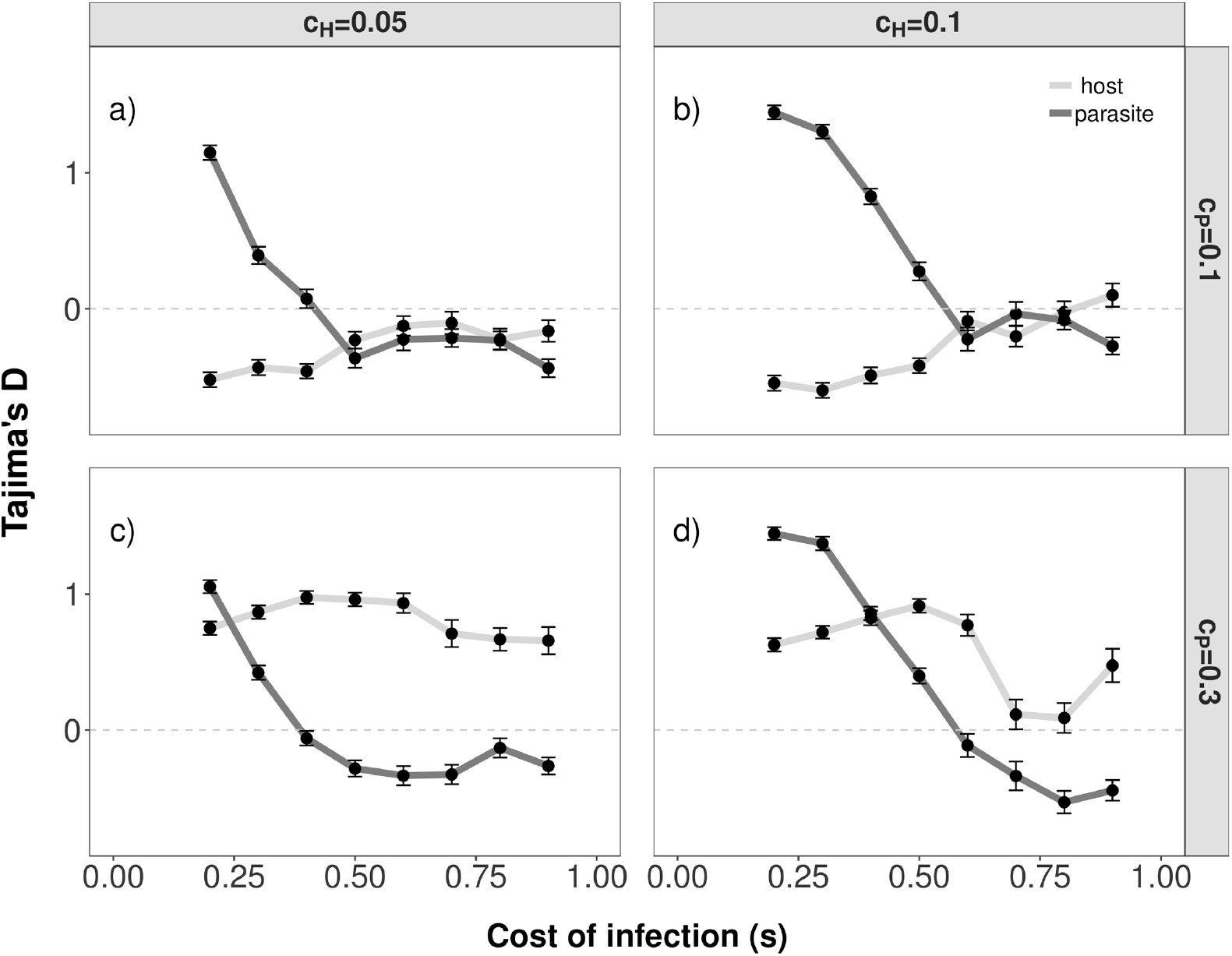
Tajima’s D model A for different costs of infection, resistance and infectivity. Tajima’s D (y-axis) for model A for various cost of infection *s* (x-axis). The results are shown for different combinations of *c*_*P*_ (*c*_*P*_ = 0.1 top, *c*_*P*_ = 0.3 bottom) and *c*_*H*_ (*c*_*H*_ = 0.05 left, *c*_*H*_ = 0.1 right). The mean and standard error of Tajima’s D of the parasite population (dark grey) and of the host population (light grey) are plotted for *r* = 200 repetitions. The dashed-dotted line shows the expected value of Tajima’s D in a Wright-Fisher population with constant population size. Tajima’s << 0 is an indicator of selective sweeps Tajima’s D >> 0 is an indicator of balancing selection. The other parameters are fixed to: *N*_*H*_ = *N*_*P*_ = 10,000, *n*_*H*_ = *n*_*P*_ = 50, *θ*_*H*_ = *θ*_*P*_ = 5, *µ*_*Rtor*_ = *µ*_*rtoR*_ = *µ*_*ntoI*_ = *µ*_*Iton*_ = 10^−5^.

**S10 Fig.**
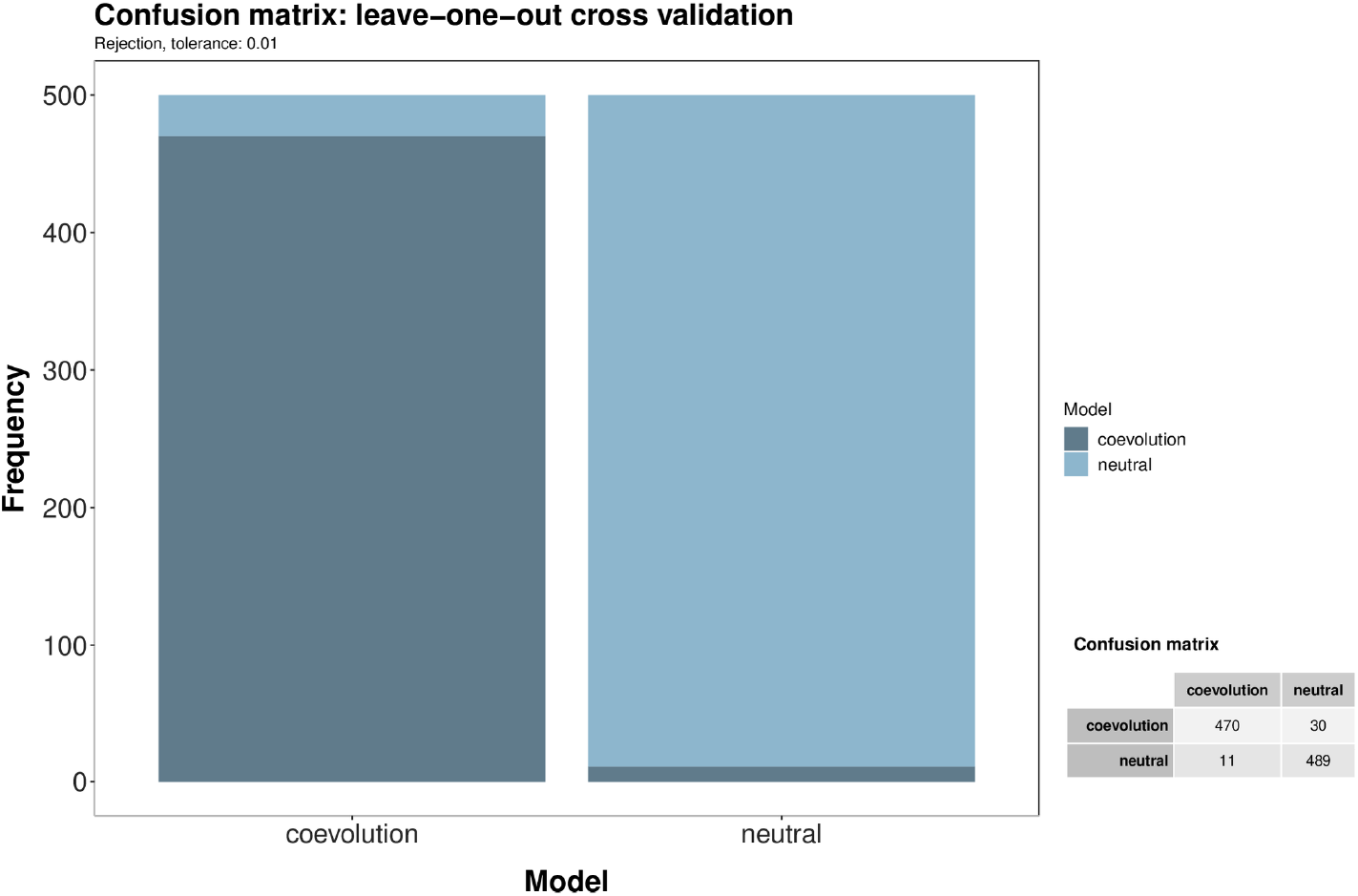
S10 Fig. Cross-validation model choice scenario 2 for r = 30 repetitions. Leave-on-out cross-validation result for distinguishing the coevolution model with unknown costs of infection (*s*), cost of resistance (*c*_*H*_) and cost of infectivity (*N*_*P*_) from a neutral model constant host and parasite population sizes (*N*_*H*_ = *N*_*P*_ = 10,000). Cross-validation results are shown for *r* = 30 and are based on 500 randomly chosen ABC-simulations for each model.

**S11 Fig.**
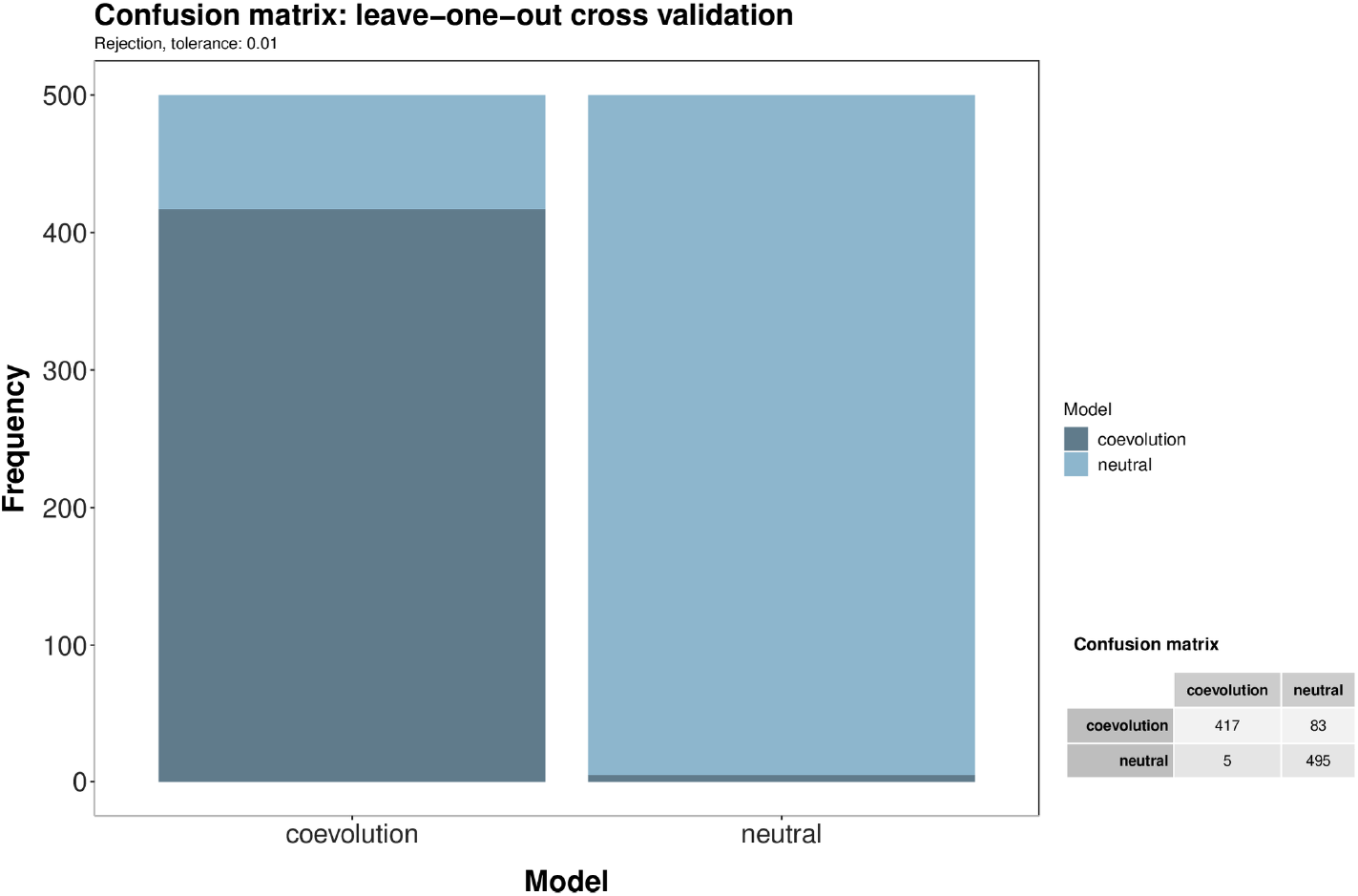
Cross-validation model choice scenario 2 for r = 10 repetitions. Leave-on-out cross-validation result for distinguishing the coevolution model with unknown costs of infection (*s*), cost of resistance (*c*_*H*_) and cost of infectivity (*N*_*P*_) from a neutral model constant host and parasite population sizes (*N*_*H*_ = *N*_*P*_ = 10,000). Cross-validation results are shown for *r* = 10 and are based on 500 randomly chosen ABC-simulations for each model.

**S12 Fig.**
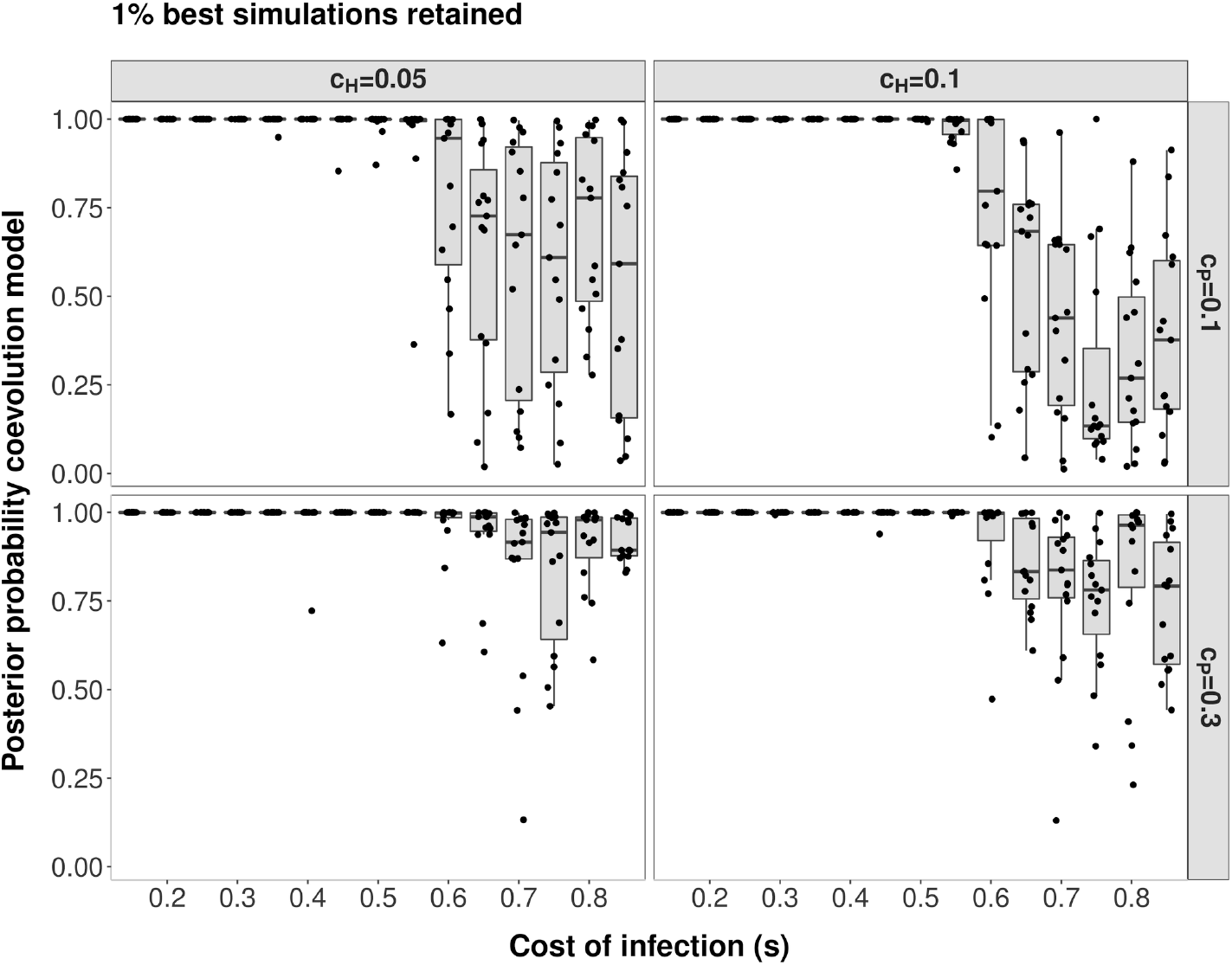
Posterior probability in support of the coevolution model (against a neutral model) for scenario 2. Results are shown for *r* = 10 and 15 PODs per boxplot. The posterior density in support of the coevolution model (y-axis) is shown for PODs with varying cost of infection (*s*). The different panels reflect the combination of *c*_*H*_ and *c*_*P*_ for the respective PODs (left: *c*_*H*_ = 0.05, right: *c*_*H*_ = 0.1, top: *c*_*P*_ = 0.1, bottom: *c*_*P*_ = 0.3). Model choice has been run to distinguish a coevolution model with unknown costs of infection (*s*), cost of resistance (*c*_*H*_) and cost of infectivity (*c*_*P*_) from a neutral model with constant host and parasite population size (*N*_*H*_ = *N*_*P*_ = 10,000). Results for single PODs are shown as dots and jitter added to the x-values to increase the readability.

**S13 Fig.**
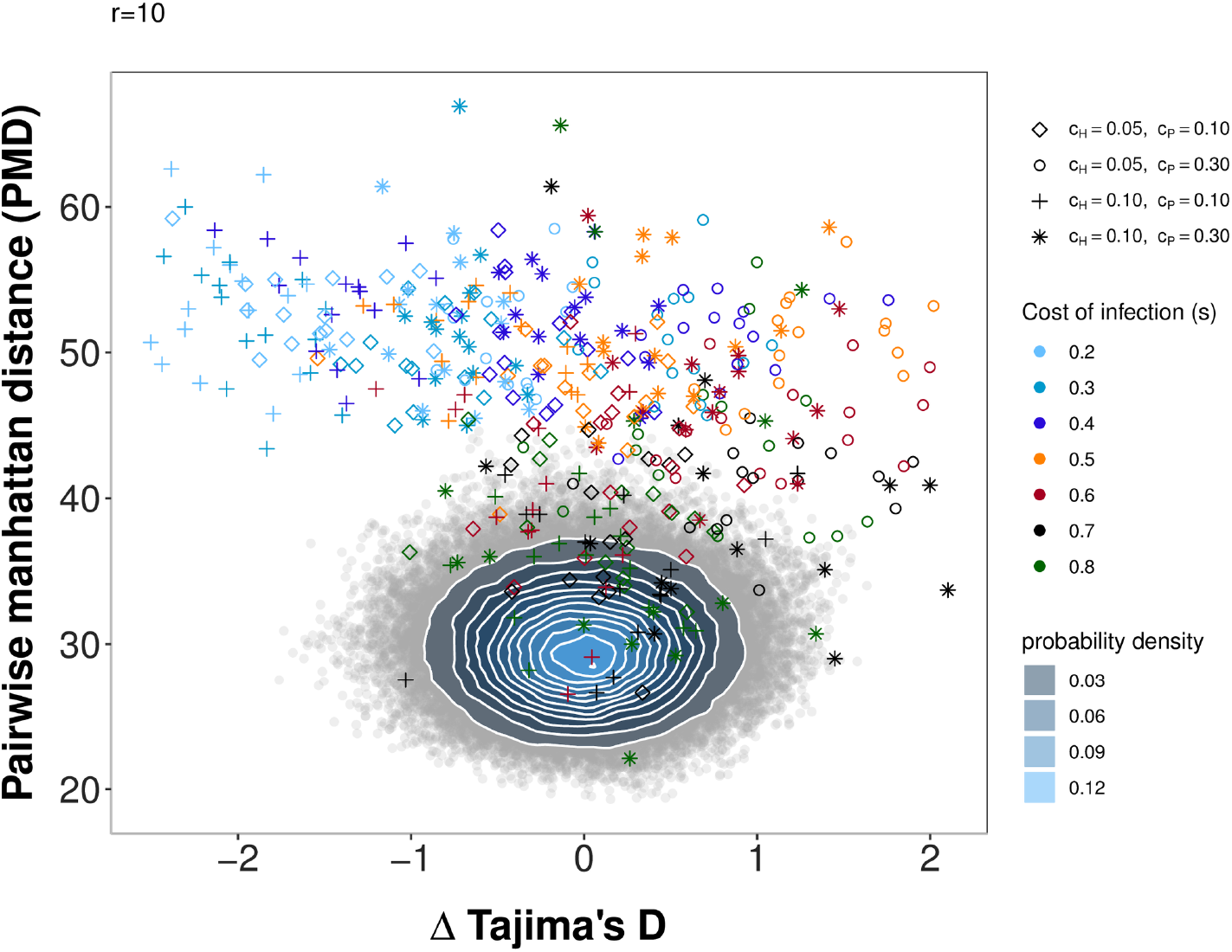
Pairwise Manhattan distance and Δ Tajima’s D (host-parasite) for the PODs under scenario 2 compared to simulations under a neutral model for r = 10. Pairwise Manhattan distance (x-axis) and the difference between Tajima’s D of the host and of the parasite (y-axis) for the PODs used for inference in Scenario 2 and 100,000 neutral simulations. Simulations under the neutral model are shown as grey open circles. A bivariate normal kernel estimation has been applied to obtain a probability density of the different summary statistic combinations. The PODs for scenario 2 are shown in color. Colors reflect the true cost of infection (*s*) for a particular POD (see legend) and shapes indicate the combination of *c*_*H*_ and *c*_*P*_ (diamonds: *c*_*H*_ = 0.05, *c*_*P*_ = 0.1; circles: *c*_*H*_ = 0.05, *c*_*P*_ = 0.3; crosses: *c*_*H*_ = 0.01, *c*_*P*_ = 0.1; stars: *c*_*H*_ = 0.1, *c*_*P*_ = 0.3) for the respective POD.

**S14 Fig.**
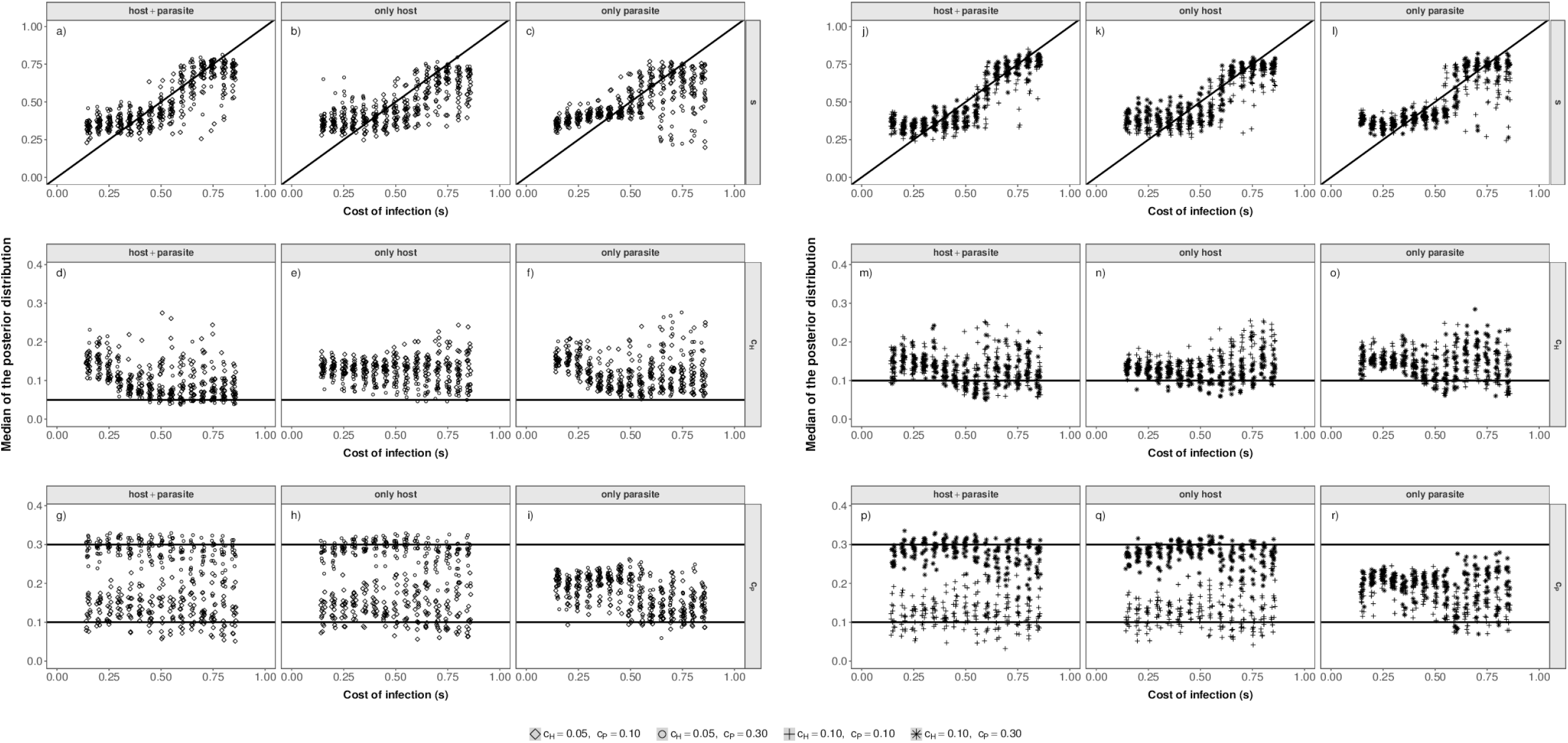
Inference results Scenario 2 for r = 10. Median of the posterior distribution (y-axis) for the cost of infection *s* (top, a-c), cost of resistance (*c*_*H*_) (middle, d-f) and cost of infectivity (*c*_*P*_) (bottom, g-i) when inference is based on host and parasite summary statistics (left), only host summary statistics (middle) or only parasite summary statistics (right) for scenario 2. The median of the posterior distribution (after post-rejection adjustment) is plotted for each POD in scenario 2. The true cost of infection for each POD is shown on the x-axis.

**S15 Fig.**
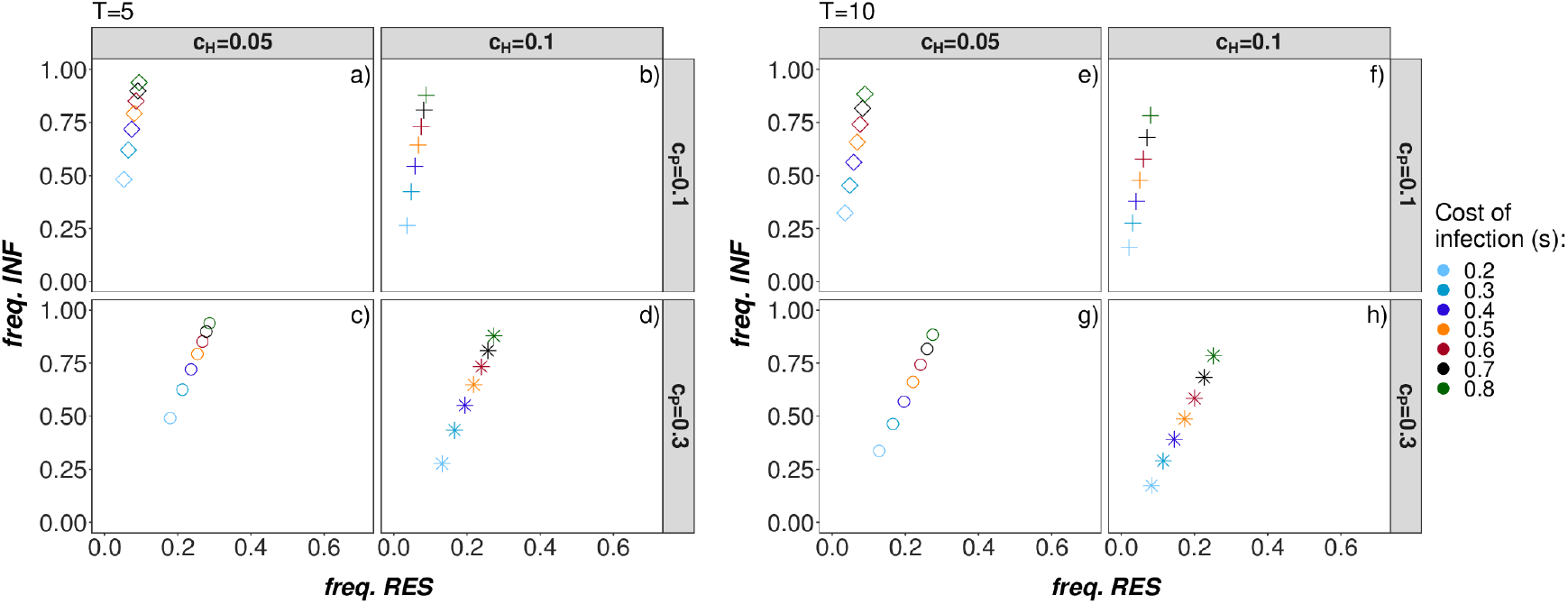
Equilibrium frequencies Model B. Deterministic equilibrium frequencies for Model B for a) *T* = 5 parasite generations (left) and b) *T* = 10 parasite generations (right) per host generation. The equilibrium frequencies for different combinations of cost of resistance *c_H_* = (0.05, 0.1) (columns), cost of infectivity *c*_*P*_ = (0.1, 0. 3) (rows) and cost of infection *s* = (0.2, 0.3, 0.4, 0.5, 0.6, 0.7, 0.8) (color of the squares) are shown. Only combinations with trench-warfare dynamics are shown. Centres of the squares represent the equilbrium frequencies obtained by simulating numerically the recursion equations in S1 File for *g*_*max*_ = 30,000 host generations starting with an initial frequency of *R*_0_ = 0.2 resistant hosts and *a*_0_ = 0.2 infective parasites.

**S16 Fig.**
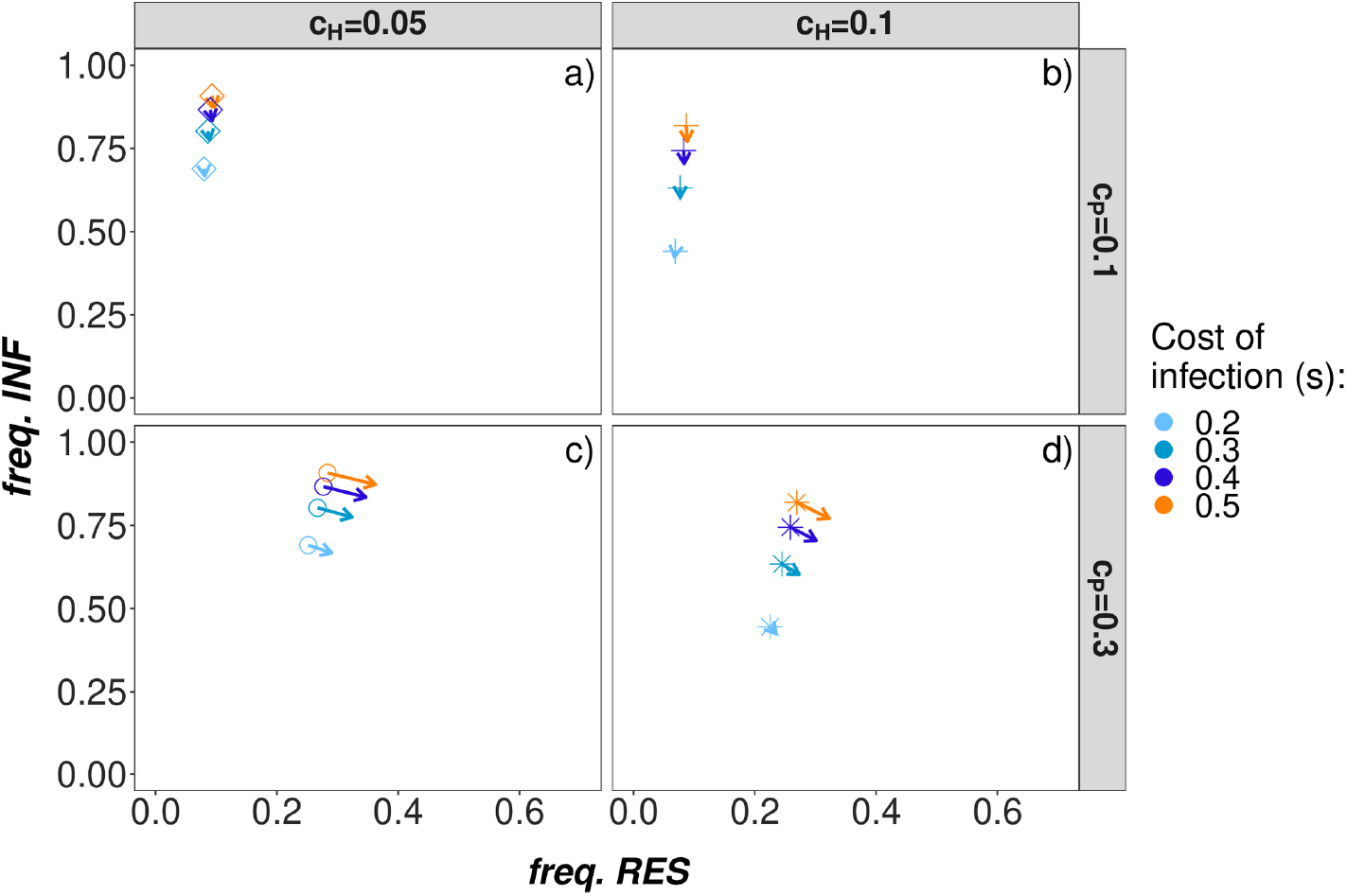
Equilibrium frequencies model C. Deterministic equilibrium frequencies for **Model C** (auto-allo-infection model) with *T* = 2 parasite generations per host generation and *ψ* = 0.95. The equilibrium frequencies for different combinations of cost of resistance *c*_*H*_ = (0.05, 0.1) (columns), cost of infectivity *c*_*P*_ = (0.1, 0. 3) (rows) and cost of infection *s* = (0.2, 0.3, 0.4, 0.5, 0.6, 0.7, 0.8) (color of the squares) are shown. Only combinations which result in trench-warfare dynamics are plotted. Centres of the squares represent the equilbrium frequencies obtained by simulating numerically the recursion equations in S1 File for *g*_*max*_ = 30,000 host generations starting with an initial frequency of *R*_0_ = 0.2 resistant hosts and *a*_0_ = 0.2 infective parasites. Heads of the arrows represent the equilibrium frequencies based on Eq (3) which corresponds to the case *ψ* = 1 [24].

**S17 Fig.**
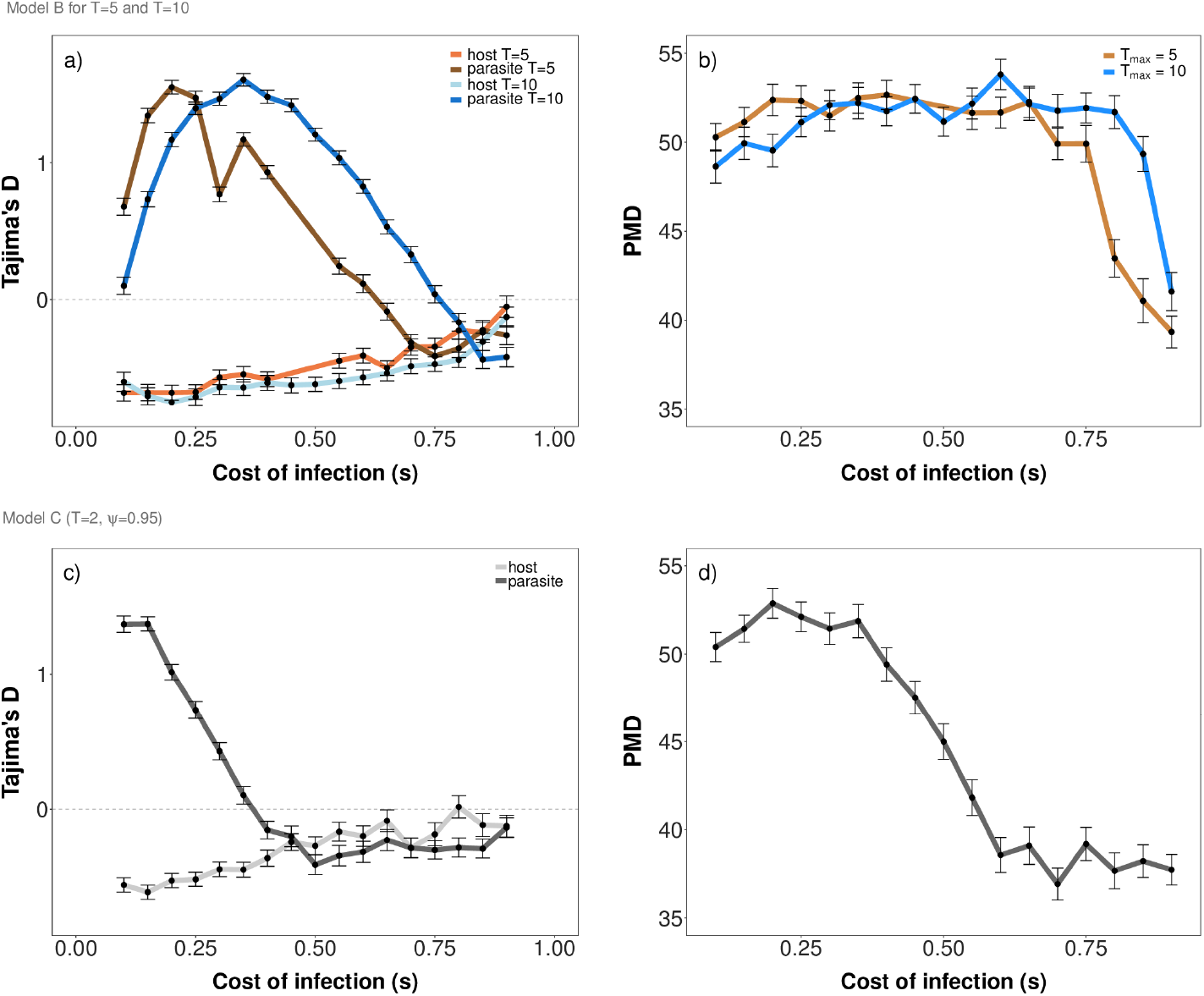
Tajima’s D and pairwise manhattan distance Model B and C. Mean and standard error of Tajima’s D (a+c) and pairwise manhattan distance (PMD) (b+d) for various costs of infection *s* (x-axis) and *r* = 200 repetitions. Results for **Model B** (pure autoinfection model with *T* = 5 and *T* = 10) are shown at the top, results for **Model C** (auto-allo-infection model with *ψ* = 0.95) are shown at the bottom. The other parameters are fixed to: *c*_*H*_ = 0.05 and *c*_*P*_ = 0.1. Initial frequencies *R*_0_ and *a*_0_ in *a* and *b* are chosen randomly from a uniform distribution between 0 and 1 while *R*_0_ = *a*_0_ = 0.2 in *c* and *d*.

**S1 File. Additional information on coevolutionary models.**

**S2 File. Details Pairwise Manhattan Distance (PMD).**

